# Non-invasive Fluorescence Imaging of Gut Commensal Bacteria in Live Mice

**DOI:** 10.1101/2022.05.05.489569

**Authors:** Alexis J. Apostolos, Mahendra D. Chordia, Sree H. Kolli, Melanie R. Rutkowski, Marcos M. Pires

## Abstract

In mammals, gut commensal microbiota interact extensively with the host and the same interactions can be dysregulated in diseased states. The development of methods to monitor gut microbiota *in vivo* can lead to improved foundational understanding of the biological events underpinning these interactions. The current standard for non-invasive monitoring of gut bacteria entails classification by 16*S* rRNA sequencing from fecal samples. This method has many advantages but also has serious limitations, especially for monitoring dynamic changes in the gut of live animals. In recent years, several imaging techniques have been widely adopted that afford non-invasive assessment of animal subjects – most notably in cancer biology; however, these technical gains have not translated to the imaging of gut bacterial communities. Herein, we describe a method to non-invasively image commensal bacteria based on the specific metabolic labeling of bacterial cell walls to illuminate the gut bacteria of live mice. This tagging strategy may additionally provide unprecedented insight into cell wall turnover of gut commensals, which has implications for bacterial cellular growth and division, in a live animal.

## Introduction

Molecular imaging of animal models is a powerful technique that is widely utilized to diagnose, measure, and track biological processes and their functional implications. Yet, to date, there are no existing standard techniques to image gut bacteria in live animals. There is a critical need for such methods as trillions of bacteria reside within the gastrointestinal (GI) tract of humans.^1, 2^ The importance of the gut community to the host is evidenced by the wide range of pathologies linked to microbiota disruptions. These range from immune dysfunctions (e.g., allergies and numerous autoimmune conditions) to endocrine system disorders (e.g., obesity and diabetes).^2-5^ Very recently, fragments of peptidoglycan (PG) within the bacterial cell wall were shown to play a major role in mediating gut-brain communication by regulating appetite and body temperature in mice^6^ and in strongly modulating responses to check-point inhibitors of immunotherapies in mouse tumor models.^7^ In the case of severe disruptions to the gut bacterial communities (e.g., colitis caused by *Clostridium difficile*), it is more straight forward to establish a clear connection and causational link.^8, 9^ But, more often, disruptions to gut commensals are subtle and chronic, requiring a range of techniques to adequately decipher them.

The current state-of-the-art, non-invasive technique used to monitor gut bacteria is classification by 16*S* rRNA sequencing from fecal samples.^10^ This method has several advantages (relatively high convenience, compositional resolution, and prolonged analysis over time) but it also has serious limitations (time gap between collection and introduction of variable, live dynamic bacterial composition of the gut may not be represented in the fecal sample, and lack of spatial resolution). Alternatively, non-invasive metabolic molecular imaging could report on dynamic responses to biological factors (e.g., prebiotics, probiotics, diet, and antibiotics) in real-time. Surprisingly, live animal imaging of gut bacterial communities remains underdeveloped, despite the growing evidence that gut bacteria play central roles in human health and disease. To directly address this technical gap, we describe a novel, non-invasive imaging modality to illuminate gut bacteria in live mice using near infrared (NIR) fluorophores by harnessing the ability of bacteria to metabolically incorporate exogenous cell wall analogs (**Figure 1A**).

**Figure 1.**
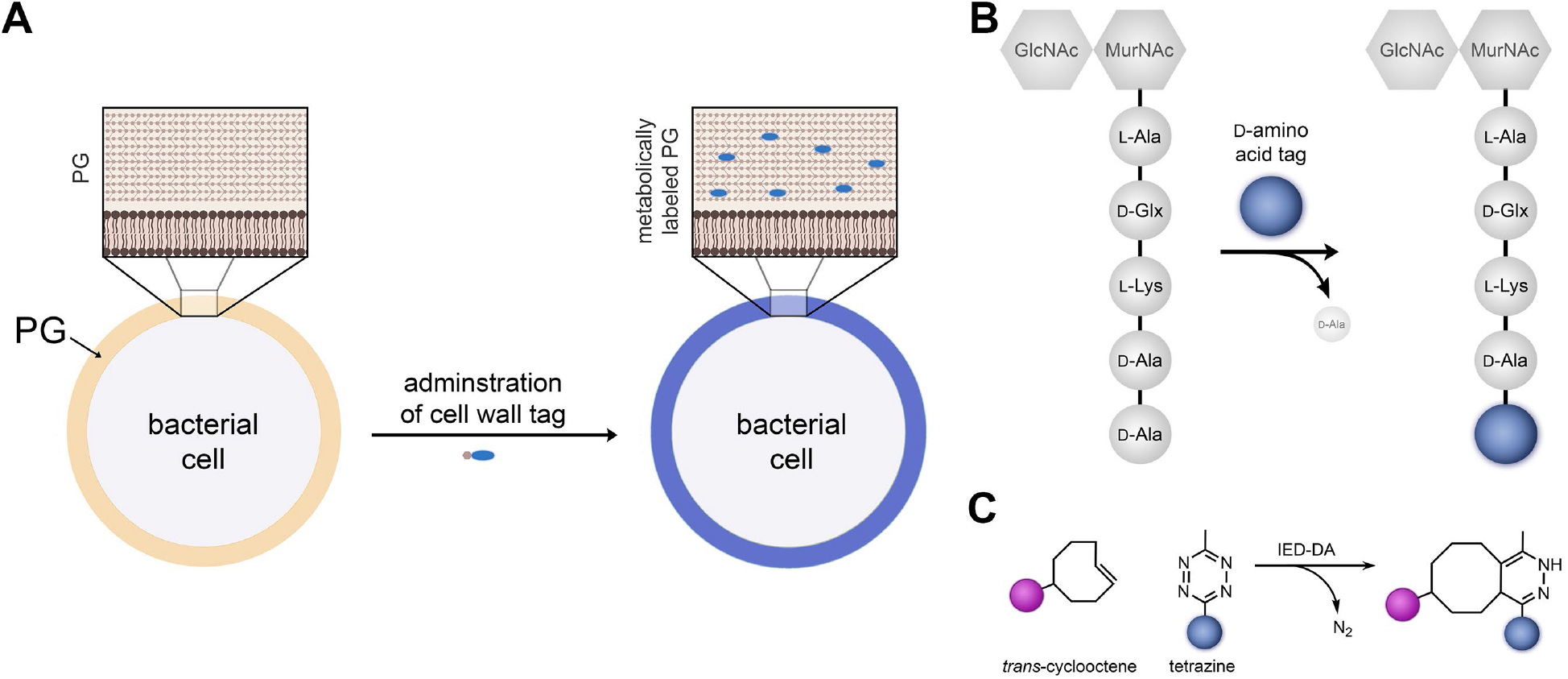
(**A**) Schematic representation of the installation of molecular fragments linked to NIR fluorescent tags within the cell wall of bacteria. (**B**) Schematic representation of an example of a PG building block in which an exogenous D-amino acid can be exchanged with D-alanine within the PG of live bacteria. (**C**) Representation of the two reactants and the product of IED-DA reactions.

Nearly all bacteria, including those residing in the gut of humans, are protected by a shield-like structure called the bacterial cell wall.^11-13^ The unique chemical composition and dynamic remodeling of the bacterial cell wall makes it an ideal target for designing metabolic molecular imaging agents with high specificity because mammalian cells do not have an equivalent biomacromolecule. A principal component of the cell wall is PG, a polymer composed of repetitive disaccharide units (*N-*acetylglucosamine and *N-* acetylmuramic acid) with short peptides (called stem peptides) linked to the *N-* acetylmuramic acid (**Figure 1B**). Neighboring stem peptides are heavily crosslinked by cell wall-linked transpeptidases to endow greater rigidity and strength to the PG matrix.^11, 14, 15^ One of the unique structural characteristics of PG is the inclusion of D-amino acids within the stem peptide.^11, 16^ This unique feature has been leveraged for the development of PG-specific labels. More specifically, bacteria can swap exogenous D-amino acids from the medium into their expanding PG scaffold during cellular growth *via* surface bound transpeptidases (**Figure 1B**).^17-27^

We posited that live gut bacteria imaging can be achieved in mice by oral administration of D-amino acid surface tags to enable the selective installation of NIR fluorophores. In 2017, Kasper et al. (and since then others^28-30^) used a D-amino acid conjugated to a visible range fluorophore to label gut bacteria of mice. These seminal results demonstrated that gut bacteria can be remodeled with single D-amino acid probes; however, visible-range fluorophores cannot be used for live animal imaging.^31^ Also in 2017, our laboratory independently demonstrated *in vivo* labeling of bacteria in the gut of *Caenorhabditis elegans* (*C. elegans*)^32^ using a 2-step strategy mediated by an inverse electron demand Diels-Alder (IED-DA) reaction.^18, 33, 34^ We showed that tetrazine-displaying D-amino acids tagged the surface of bacteria in *C. elegans* and after treatment with a *trans*-cyclooctene (TCO)-linked Cy5 fluorophore, bacterial cells were selectively labeled with the fluorophore. IED-DA reactants include TCO and tetrazine moieties, which rapidly and quantitatively react to form a stable covalent bond (**Figure 2C**).^35^ Critically, the biocompatibility of the tetrazine/TCO handles is highlighted by the first successful completion of a human clinical trial in which doxorubicin-TCO was periodically administered to selectively release drugs from a tetrazine-modified polymer implanted within tumor masses.^36^

**Figure 2.**
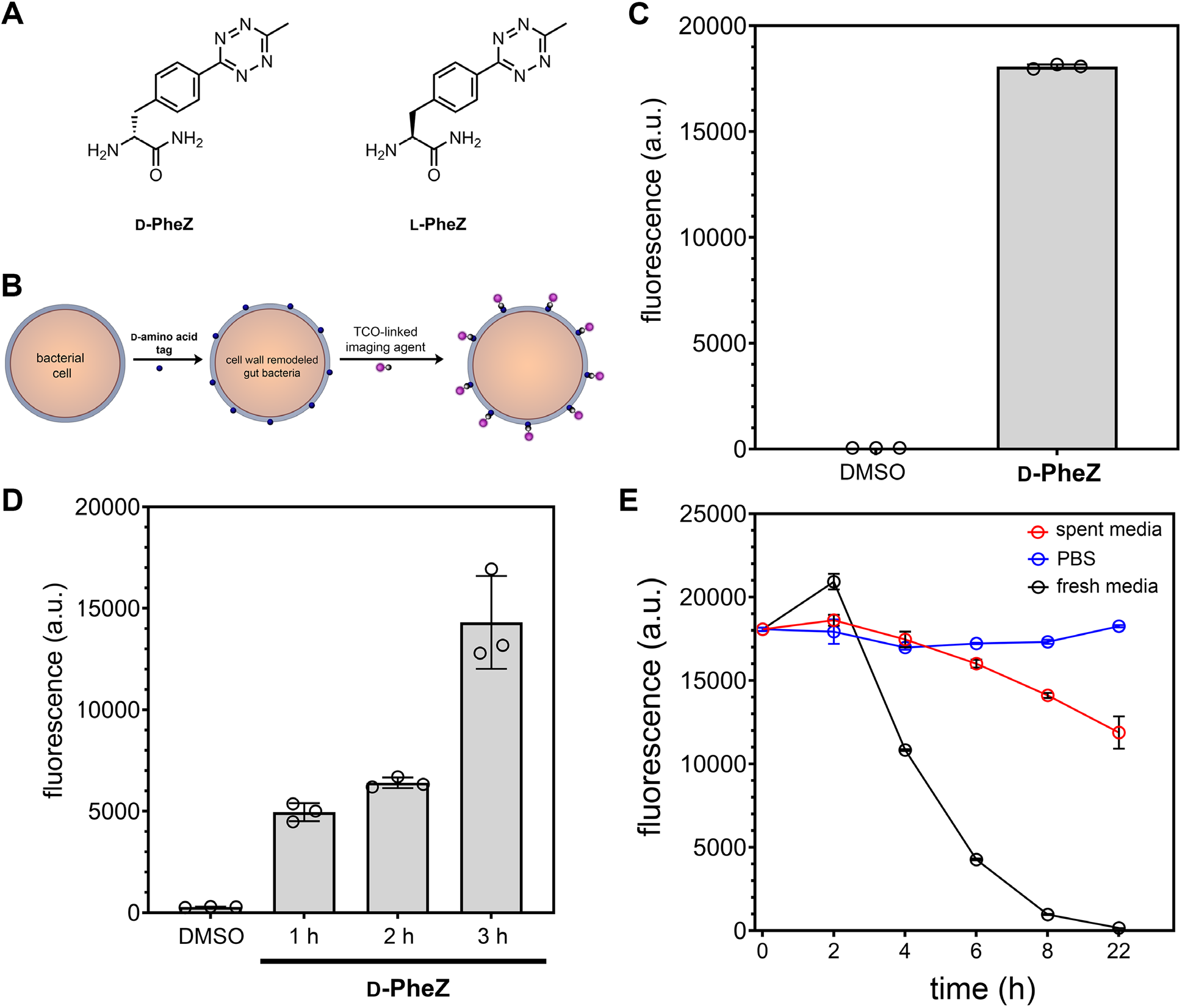
(**A**) Chemical structures of **D-PheZ** and **L-PheZ**. (**B**) Schematic representation of the steps leading to surface tagging of bacterial cells following metabolic incorporation of D-amino acids. (**C**) Flow cytometry analysis of *L. casei* grown overnight with 25 µM **D-** or **L-PheZ** followed by treatment with 25 µM TCO-Cy5. (**D**) Flow cytometry analysis of *L. casei* grown to stationary phase and then treated with 25 µM **D-PheZ**, followed by TCO-Cy5 at different time points. (**E**) Flow cytometry analysis of *L. casei* treated overnight with 100 µM of **D-PheZ**. Subsequently, cells were incubated with 50 µM of TCO-Cy5 and then resuspended in the conditions noted. Data are represented as mean +/-SD (n = 3). *P*-values were determined by a two-tailed *t*-test (* denotes a *p*-value < 0.05, ** < 0.01, ***<0.001, ns = not significant).

As a starting point in the development of the tagging strategy, we set out to adopt our earlier findings with *C. elegans* to NIR imaging. To this end, we utilized the combination of a tetrazine-modified D-amino acid (**D-PheZ**) and a TCO-linked fluorophore (**Figure 2A**). Surface tagging conditions were first analyzed *in vitro* by incubation of bacteria with the metabolic tag followed by labeling with the reactive fluorophore, which is expected to fluorescently tag the PG of live bacteria (**Figure 2B**). However, when transitioning to *in vivo* studies, a key feature to consider when developing a dosing and imaging schedule is the kinetics of PG biosynthesis and remodeling of gut bacteria. It is well established that PG biosynthesis and remodeling is tightly coupled to bacterial cell growth, which requires new PG material to be assembled to support *de novo* cellular replication.^11, 14, 15^ While these features can be readily assessed and measured *in vitro*, they remain largely uncharacterized in commensal communities. Recent efforts have started to provide estimates into the doubling rate of gut bacteria in mice.^37, 38^ However, these tools have limitations in that they primarily measure relative growth rates and they require the use of non-native, genetically modified organisms. As such, questions regarding the rate of gut bacteria PG turn-over in gut commensals are yet to be fully elucidated.

In the context of our assay, if there is considerably rapid PG biosynthesis (consistent with fast bacterial replication), the expectation would be that the PG probe would be readily incorporated into the cell wall, as it is typically observed in a nutrient-rich culture. To mimic fast-growth kinetics, *Lactobacillus casei* (*L. casei*) were diluted (1:100) and grown through log-phase in the presence of **D-PheZ**, followed by treatment with TCO-Cy5 dye (**Figure 2C**). *L. casei* is commonly found in the gut of humans and can be used a model for PG remodeling.^39^ As expected, and similar to our previous results using other bacterial species, cellular fluorescence levels were significantly higher than untreated cells. Cells treated with **L-PheZ** showed a significantly lower level of fluorescence compared to treatment with **D-PheZ**. These results are anticipated because the D-enantiomer of non-canonical amino acids has been shown to be preferentially incorporated in the PG, while modified L-amino acids are not.^17^ Alternatively, we considered the possibility that bacterial cells in the gut may be in a slower-replicating (or quiescent) state. To approximate this, **D-PheZ** was added to stationary phase *L. casei* and fluorescence levels were monitored over time (**Figure 2D**). A clear increase in cellular fluorescence was observed despite limited cellular growth. It is likely that PG remodeling, rather than PG biosynthesis, could account for this increase in metabolic tagging.

Additionally, the kinetics of loss of the metabolic tag needed to be considered. To model this, *L. casei* from an overnight culture was labeled with the combination of **D-PheZ** and TCO-Cy5. Cells were washed, the supernatant was removed and replaced with either PBS or media isolated from *L. casei* in stationary growth (spent media). A third condition involved the back-dilution of cells into fresh media. From these results, it is evident that fresh media stimulates a rapid loss of surface tagging, presumably due to faster cellular growth and PG remodeling kinetics in the absence of the PG probe. Media promoting slower growth kinetics showed considerably slower loss in cellular fluorescence (**Figure 2E**). Therefore, we anticipate that loss of cellular tagging could be a proxy for the kinetics of cellular growth and remodeling, and, therefore, may be used to gain insight into PG remodeling in gut commensal bacteria.

While Cy5 can be used to image bacteria in the transparent *C. elegans*, it was necessary to switch to a NIR fluorophore (Cy7.5) to properly image gut commensal bacteria in mice. *In vitro* labeling of bacteria was performed to establish the suitability of TCO-Cy7.5 in PG tagging. Three species of bacteria were chosen: *L. casei, Enterococcus faecalis* (*E. faecalis*), and *Lactobacillus plantarum* (*L. plantarum*). These organisms have been previously found in the gut of humans.^40^ Cells were individually subjected to the 2-step surface tagging scheme and analyzed by flow cytometry (**Figure S1**). As expected due to the structural similar of Cy7.5 to Cy5, high levels of cellular fluorescence were observed across all species of bacteria. Background cellular fluorescence levels were low and, similarly, treatment with the dye alone (TCO-Cy7.5) led to minimal cellular fluorescence. These confirmatory results led us to test a similar bacterial labeling scheme in live mice. For these experiments, mice were first orally gavaged with **D-PheZ** or **L-PheZ**, followed by a 2-hour chase period (to allow the unincorporated amino acid to passage), then administered with TCO-Cy7.5. Mice were imaged *via* NIRF-IVIS two hours following the gavaging with the dye (**Figure 3**). A measurable difference in fluorescence was observed in mice treated with the **D-PheZ** relative to the stereocontrol **L-PheZ**, a finding that is suggestive of cell wall labeling of gut commensals. Interestingly, it was observed that the labeling half-life was relatively short using **D-PheZ**, as evidenced by the decrease in fluorescence signal between 4 and 22 hours after the final dosing period (**Figure S2**). Together, these results establish that single D-amino acids can potentially be applied to illuminate the gut commensal community of live animals.

**Figure 3.**
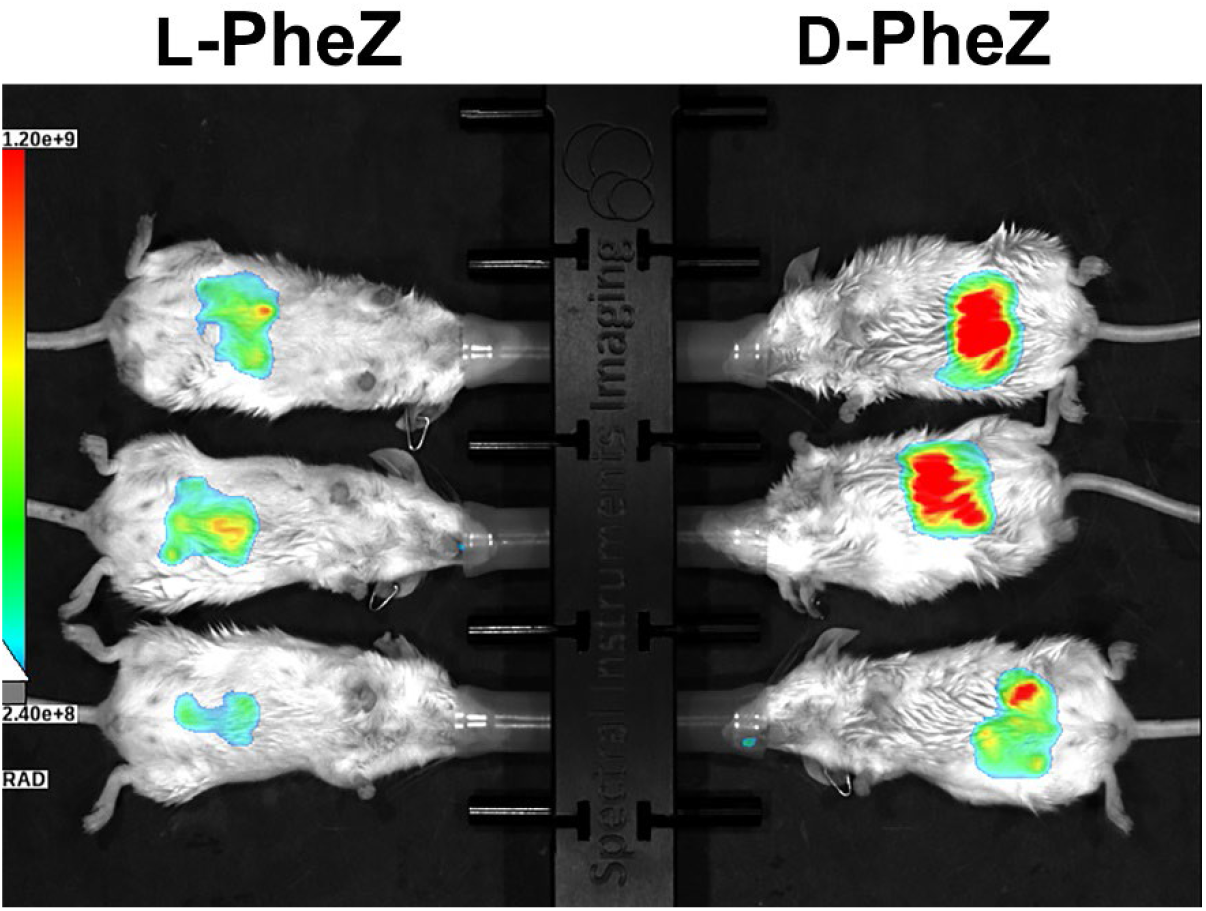
IVIS imaging of WT female mice orally gavaged with **L-PheZ** or **D-PheZ** (5 mM in 200 µL of PBS) twice (1 hour apart). After 4 hours, mice were orally administered TCO-Cy7.5 (1 mM in 100 µL of PBS). Imaging was performed 2 hours after dye administration.

Encouraged by the results of single D-amino acid gut labeling, and as an alternative strategy, we sought to image gut bacteria by using a one-step method in which the fluorophore is connected directly to the metabolic tag. A one-step labeling strategy is aimed at reducing the potential timing and cross-reactivity issues that could be associated with the 2-step IED-DA strategy. One option would be to directly conjugate the fluorophore onto the sidechain of a single D-amino acid (e.g., D-lysine). However, we hypothesized that large sidechains would significantly impede PG incorporation due to poor recognition by cell wall transpeptidases. This hypothesis was premised on our previous systematic study using a library of D-amino acid variants in which the fluorophore was held constant but the chemical structure of the sidechain varied in physical characteristics (size, length, flexibility, charge).^34^ Because NIR fluorophores are typically large (and often highly charged to compensate for the hydrophobicity of the elongated conjugated system), we had anticipated that modification of a NIR fluorophore onto the D-amino acid sidechain would greatly suppress PG incorporation. Instead, we envisioned that administration of a synthetic stem peptide analog conjugated to a NIR fluorophore would circumvent this challenge (**Figure 4A**). We^41-44^, and others^45-47^, have shown that treatment of bacterial cells with stem peptide analogs also result in their incorporation the PG by PG-related transpeptidases (**Figure 4B**). The principal advantage of stem peptide analogs is that our laboratory previously showed that the *N*-terminus of a tetra-or penta-stem peptide analog demonstrates tolerability to large chemical modifications.

**Figure 4.**
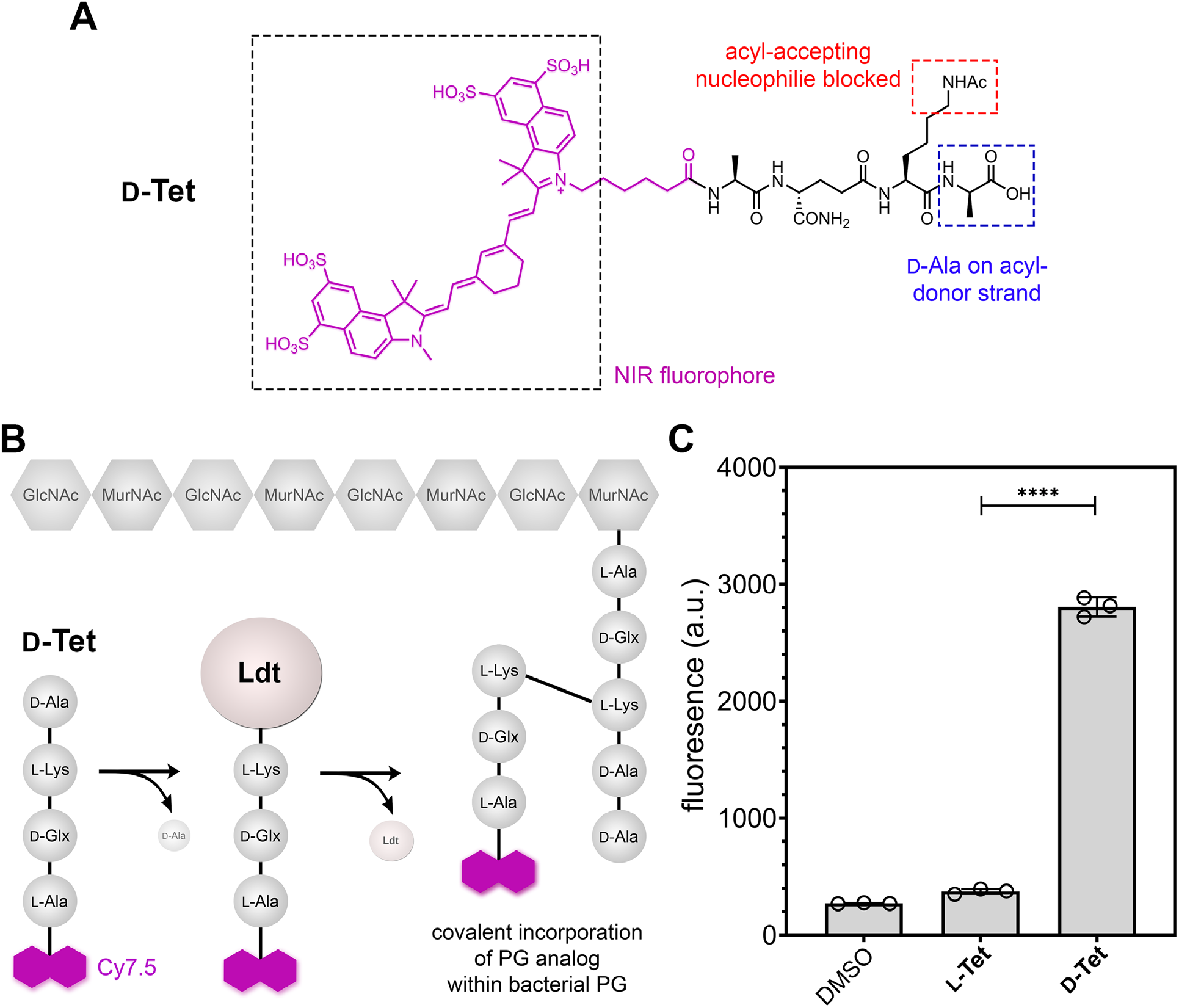
(**A**) Chemical structure of **D-Tet** and description of the three primary design features. (**B**) Schematic representation of the mode of incorporation of **D-Tet** within existing PG in live bacteria. The tetrapeptide is expected to be processed by transpeptidases by the removal of the terminal D-Ala, which yields a thioester intermediate in the active site of Ldts. Subsequent capture by a nearby nucleophile leads to the formation of a stable covalent bond between the PG and the imaging probe. (**C**) *L. casei* treated overnight with 100 µM of **L-** or **D-Tet** and subsequently analyzed by flow cytometry. Data are represented as mean +/-SD (n = 3). *P*-values were determined by a two-tailed *t*-test (* denotes a *p*-value < 0.05, ** < 0.01, ***<0.001, ns = not significant).

We synthesized two analogs of the canonical stem peptide from PG with a NIR Cy7.5 covalently linked to the *N*-terminus (**Figure S3**). One of the tetrapeptides (**D-Tet**) contained a D-alanine on the *C*-terminus, which mimics the native stem peptide of almost all known bacteria. We also synthesized a control diastereomer (**L-Tet**) in which a single stereocenter on the *C*-terminal alanine is in the L configuration, which prevents processing by bacterial transpeptidases.^48^ As a starting point, incorporation of the tetrapeptide tag (**D-Tet**) was first assessed *in vitro* (**Figure 4C**). As expected, bacterial cells treated with **D-Tet** displayed notably higher cellular fluorescence relative to untreated cells and cells treated with the diastereomer **L-Tet**. With these results, we set out to evaluate the tetra-peptide probes *in vivo*. Mice were orally gavaged either **D-Tet** or **L-Tet** and imaged periodically afterwards using NIRF imaging (**Figure 5A**). Interestingly, in early time points there was minimal difference between the two treatment groups, which is likely reflective of tagging reagents undergoing peristaltic transit through the GI tract (**Figure S4**). After a chase period, there was a measurable difference in the fluorescence levels of mice administered the cell wall tags. Histological studies of the GI tract of mice treated with **D-Tet** was similar to that of untreated mice, which underscores the selectivity of metabolic incorporation of the probe and its lack of adverse interaction with the host cells (**Figure S5**) These results, collectively, suggest that a stem-peptide analog **D-Tet** can selectively tag and report on the presence of gut bacteria in live mice.

**Figure 5.**
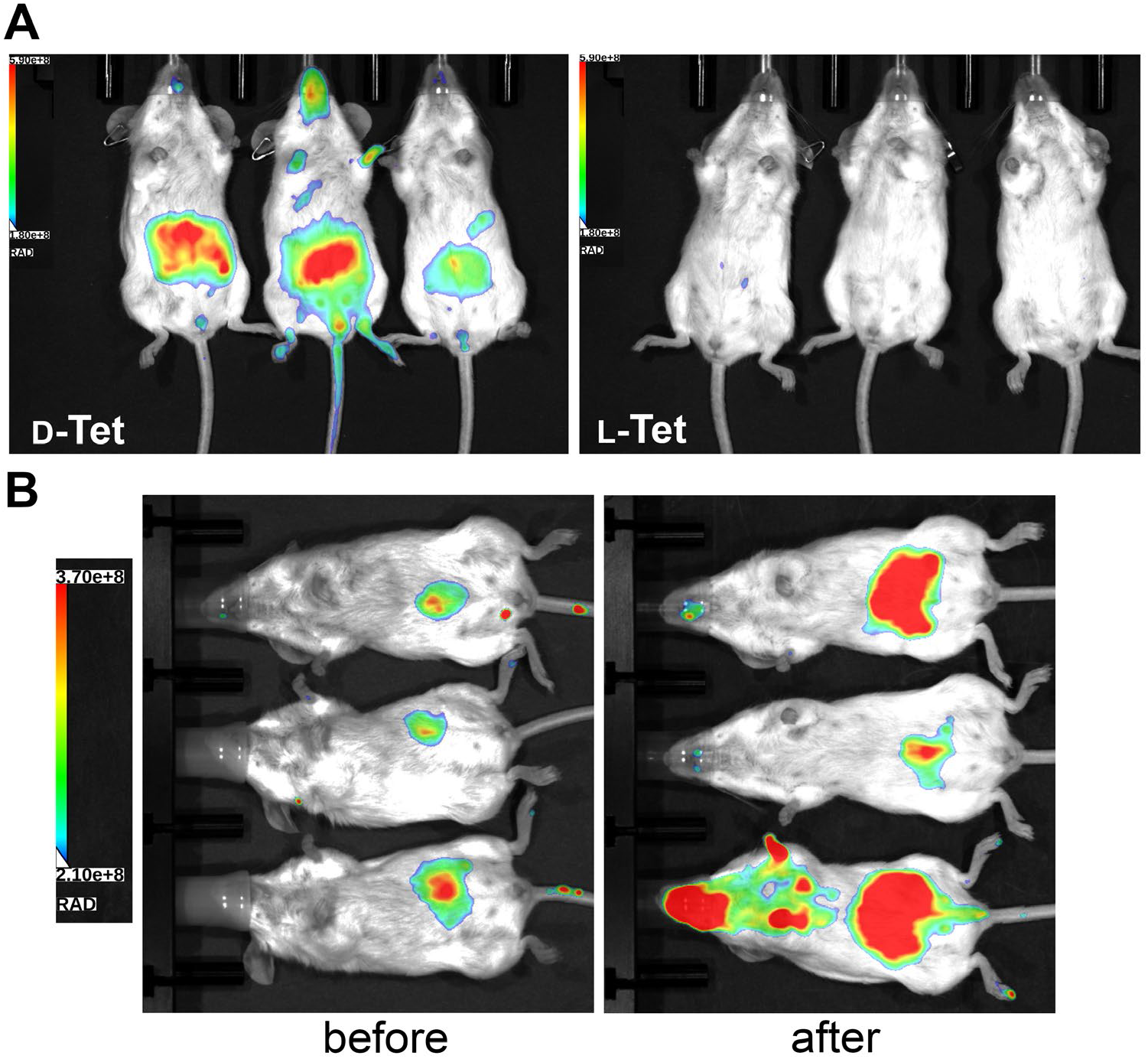
(**A**) IVIS imaging of female mice orally gavaged 2x, 1 h apart, with a tetrapeptide tag (1 mM in 100 µL of PBS) containing a terminal D-Ala (**D-Tet**), expected not to incorporate in the bacterial PG (lef*t*) and with a tetrapeptide tag containing a terminal L-Ala (**L-Tet**), expected to incorporate in the cell walls of bacteria in the mouse gut (*right*). Mice were imaged 22 hours after the final administration. (**B**) *Left*: IVIS imaging of GF BALB/cAnNTac mice orally gavaged 2x, 1 h apart, with **D-Tet** tag (1 mM in 100 µL of PBS) before conventionalization and imaged 22 hours after the final administration. *Right*: IVIS imaging of the same mice after conventionalization. Mice were orally gavaged 2x, 1 h apart, with **D-Tet** tag (1 mM in 100 µL of PBS) and imaged 22 hours after the final administration. On the day of the gavage, one of the mice bit through the galvaging tube resulting in evident staining of the mouth and paw areas, which is visible in the left panel of 6B (bottom mouse).

To further evaluate the utility and applicability of this labeling strategy, we sought to test the imaging of Germ-Free (GF) mice (BALB/cAnNTac) that are devoid of a gut bacterial community. The lack of bacterial cells that can be metabolically labelled by our probe is expected to result in background levels of signals. These mice were gavaged with **D-Tet** as before and imaged (**Figure 5B**). The same mice were subsequently conventionalized by co-housing with SPF mice for 4 days to allow for a period of recovery upon introduction to a conventional vivarium. This was followed by a series of fecal microbiota transplantation (FMT) treatments (3 times, once a day) to robustly induce bacterial engraftment in the recipient mice. The mice were co-housed for an additional 10 days to promote the growth and repopulation of the gut bacterial communities. All together, these measures are consistent with the amount of time that is required to observe a stable level of bacterial cells in the GI tract of GF mice post FMT from donor SPF mice.^49, 50^ Oral gavaging of **D-Tet** to mice was performed 2X on day one as described previously, followed by NIR imaging analysis 22 hours later. Our results show a marked increase in fluorescence (**Figure 5B**) in mice that were conventionalized relative to GF mice, which is consistent with the expected engraftment of bacterial cells. *Ex-vivo* imaging of the GI tract of the same mice demonstrated that the cecum and the large intestines appear to label more prominently, which would be consistent with a larger load of bacterial cells (**Figure S6**). These results represent a promising first step to potentially extend this strategy in the future into a non-invasive imaging tool that can provide insight into the dynamics of the gut commensal bacteria.

In conclusion, we have demonstrated that cell wall probes can provide a potential route to the non-invasive imaging of gut commensal bacteria in mice. To this end, animals were administered synthetic analogs of PG (either single D-amino acids or tetrapeptide stem mimetics) to induce their metabolic incorporation into the PG scaffold. Single amino acid labeling was based on biorthogonal installation of a NIR fluorophore while the tetrapeptide was directly linked to the NIR fluorophore. Overall, there was discernable differences observed in both routes of tagging; however, the tetrapeptide labeling showed a stronger contrast between the metabolic tag and a stereocontrol. Interestingly, there was a relatively short half-life of the single D-amino acid PG label, which may indicate that the cell walls of gut commensals undergo considerable levels of PG remodeling. Finally, we demonstrated that our labeling strategy could illuminate the colonization of GF mice following FMT. In the future, we plan to optimize the labeling conditions to gain greater insight into the dynamics related to cell wall biology in gut bacteria of live mice in response to various stressors (antibiotics, bacteriophage, and probiotics).

## Supporting Information

Additional figures, tables, and materials/methods are included in the supporting information file.

## Table of Content Figure

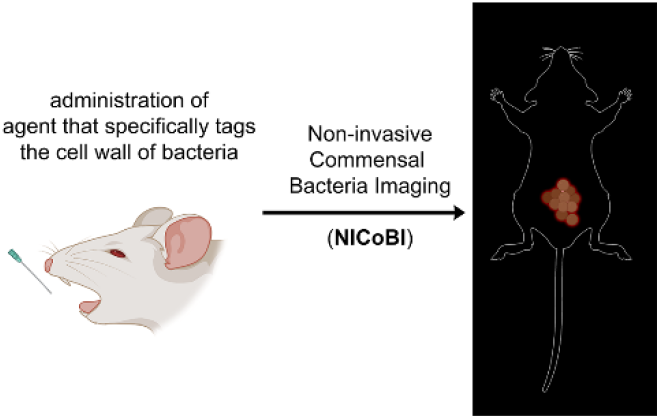

## References

1. Sender, R.; Fuchs, S.; Milo, R., Revised Estimates for the Number of Human and Bacteria Cells in the Body. PLoS Biol 2016, 14 (8), e1002533.

2. Fan, Y.; Pedersen, O., Gut microbiota in human metabolic health and disease. Nat Rev Microbiol 2021, 19 (1), 55–71.

3. Shreiner, A. B.; Kao, J. Y.; Young, V. B., The gut microbiome in health and in disease. Curr Opin Gastroenterol 2015, 31 (1), 69–75.

4. Guarner, F.; Malagelada, J. R., Gut flora in health and disease. Lancet 2003, 361 (9356), 512–9.

5. Belkaid, Y.; Hand, T. W., Role of the microbiota in immunity and inflammation. Cell 2014, 157 (1), 121–41.

6. Gabanyi, I.; Lepousez, G.; Wheeler, R.; Vieites-Prado, A.; Nissant, A.; Wagner, S.; Moigneu, C.; Dulauroy, S.; Hicham, S.; Polomack, B.; Verny, F.; Rosenstiel, P.; Renier, N.; Boneca, I. G.; Eberl, G.; Lledo, P. M., Bacterial sensing via neuronal Nod2 regulates appetite and body temperature. Science 2022, 376 (6590), eabj3986.

7. Griffin, M. E.; Espinosa, J.; Becker, J. L.; Luo, J. D.; Carroll, T. S.; Jha, J. K.; Fanger, G. R.; Hang, H. C., Enterococcus peptidoglycan remodeling promotes checkpoint inhibitor cancer immunotherapy. Science 2021, 373 (6558), 1040–1046.

8. Baumler, A. J.; Sperandio, V., Interactions between the microbiota and pathogenic bacteria in the gut. Nature 2016, 535 (7610), 85–93.

9. Theriot, C. M.; Koenigsknecht, M. J.; Carlson, P. E., Jr.; Hatton, G. E.; Nelson, M.; Li, B.; Huffnagle, G. B.; J, Z. L.; Young, V. B., Antibiotic-induced shifts in the mouse gut microbiome and metabolome increase susceptibility to Clostridium difficile infection. Nat Commun 2014, 5, 3114.

10. Gill, S. R.; Pop, M.; Deboy, R. T.; Eckburg, P. B.; Turnbaugh, P. J.; Samuel, S.; Gordon, J. I.; Relman, D. A.; Fraser-Liggett, C. M.; Nelson, K. E., Metagenomic analysis of the human distal gut microbiome. Science 2006, 312 (5778), 1355–9.

11. Vollmer, W.; Blanot, D.; de Pedro, M. A., Peptidoglycan structure and architecture. FEMS Microbiol Rev 2008, 32 (2), 149–67.

12. Vollmer, W.; Blanot, D.; De Pedro, M. A., Peptidoglycan structure and architecture. FEMS Microbiology Reviews 2008, 32 (2), 149–167.

13. Typas, A.; Banzhaf, M.; Gross, C. A.; Vollmer, W., From the regulation of peptidoglycan synthesis to bacterial growth and morphology. Nat Rev Microbiol 2011, 10 (2), 123–36.

14. Holtje, J. V., Growth of the stress-bearing and shape-maintaining murein sacculus of Escherichia coli. Microbiol Mol Biol Rev 1998, 62 (1), 181–203.

15. Lovering, A. L.; Safadi, S. S.; Strynadka, N. C., Structural perspective of peptidoglycan biosynthesis and assembly. Annu Rev Biochem 2012, 81, 451–78.

16. Silhavy, T. J.; Kahne, D.; Walker, S., The bacterial cell envelope. Cold Spring Harb Perspect Biol 2010, 2 (5), a000414.

17. Kuru, E.; Hughes, H. V.; Brown, P. J.; Hall, E.; Tekkam, S.; Cava, F.; de Pedro, M. A.; Brun, Y. V.; VanNieuwenhze, M. S., In Situ probing of newly synthesized peptidoglycan in live bacteria with fluorescent D-amino acids. Angew Chem Int Ed Engl 2012, 51 (50), 12519–23.

18. Siegrist, M. S.; Whiteside, S.; Jewett, J. C.; Aditham, A.; Cava, F.; Bertozzi, C. R., (D)-Amino acid chemical reporters reveal peptidoglycan dynamics of an intracellular pathogen. ACS Chem Biol 2013, 8 (3), 500–5.

19. Bisson-Filho, A. W.; Hsu, Y. P.; Squyres, G. R.; Kuru, E.; Wu, F.; Jukes, C.; Sun, Y.; Dekker, C.; Holden, S.; VanNieuwenhze, M. S.; Brun, Y. V.; Garner, E. C., Treadmilling by FtsZ filaments drives peptidoglycan synthesis and bacterial cell division. Science 2017, 355 (6326), 739–743.

20. Pilhofer, M.; Aistleitner, K.; Biboy, J.; Gray, J.; Kuru, E.; Hall, E.; Brun, Y. V.; VanNieuwenhze, M. S.; Vollmer, W.; Horn, M.; Jensen, G. J., Discovery of chlamydial peptidoglycan reveals bacteria with murein sacculi but without FtsZ. Nat Commun 2013, 4, 2856.

21. Fleurie, A.; Lesterlin, C.; Manuse, S.; Zhao, C.; Cluzel, C.; Lavergne, J. P.; Franz-Wachtel, M.; Macek, B.; Combet, C.; Kuru, E.; VanNieuwenhze, M. S.; Brun, Y. V.; Sherratt, D.; Grangeasse, C., MapZ marks the division sites and positions FtsZ rings in Streptococcus pneumoniae. Nature 2014, 516 (7530), 259–62.

22. Liechti, G. W.; Kuru, E.; Hall, E.; Kalinda, A.; Brun, Y. V.; VanNieuwenhze, M.; Maurelli, A. T., A new metabolic cell-wall labelling method reveals peptidoglycan in Chlamydia trachomatis. Nature 2014, 506 (7489), 507–10.

23. Faure, L. M.; Fiche, J. B.; Espinosa, L.; Ducret, A.; Anantharaman, V.; Luciano, J.; Lhospice, S.; Islam, S. T.; Treguier, J.; Sotes, M.; Kuru, E.; Van Nieuwenhze, M. S.; Brun, Y. V.; Theodoly, O.; Aravind, L.; Nollmann, M.; Mignot, T., The mechanism of force transmission at bacterial focal adhesion complexes. Nature 2016, 539 (7630), 530–535.

24. Qiao, Y.; Lebar, M. D.; Schirner, K.; Schaefer, K.; Tsukamoto, H.; Kahne, D.; Walker, S., Detection of lipid-linked peptidoglycan precursors by exploiting an unexpected transpeptidase reaction. J Am Chem Soc 2014, 136 (42), 14678–81.

25. Lebar, M. D.; May, J. M.; Meeske, A. J.; Leiman, S. A.; Lupoli, T. J.; Tsukamoto, H.; Losick, R.; Rudner, D. Z.; Walker, S.; Kahne, D., Reconstitution of peptidoglycan cross-linking leads to improved fluorescent probes of cell wall synthesis. J Am Chem Soc 2014, 136 (31), 10874–7.

26. Shieh, P.; Siegrist, M. S.; Cullen, A. J.; Bertozzi, C. R., Imaging bacterial peptidoglycan with near-infrared fluorogenic azide probes. Proc Natl Acad Sci U S A 2014, 111 (15), 5456–61.

27. Ngo, J. T.; Adams, S. R.; Deerinck, T. J.; Boassa, D.; Rodriguez-Rivera, F.; Palida, S. F.; Bertozzi, C. R.; Ellisman, M. H.; Tsien, R. Y., Click-EM for imaging metabolically tagged nonprotein biomolecules. Nat Chem Biol 2016, 12 (6), 459–65.

28. Wang, W.; Lin, L.; Du, Y.; Song, Y.; Peng, X.; Chen, X.; Yang, C. J., Assessing the viability of transplanted gut microbiota by sequential tagging with D-amino acid-based metabolic probes. Nat Commun 2019, 10 (1), 1317.

29. Lin, L.; Song, J.; Du, Y.; Wu, Q.; Gao, J.; Song, Y.; Yang, C.; Wang, W., Quantification of Bacterial Metabolic Activities in the Gut by d-Amino Acid-Based In Vivo Labeling. Angew Chem Int Ed Engl 2020, 59 (29), 11923–11926.

30. Wang, W.; Yang, Q.; Du, Y.; Zhou, X.; Du, X.; Wu, Q.; Lin, L.; Song, Y.; Li, F.; Yang, C.; Tan, W., Metabolic Labeling of Peptidoglycan with NIR-II Dye Enables In Vivo Imaging of Gut Microbiota. Angew Chem Int Ed Engl 2020, 59 (7), 2628–2633.

31. Hudak, J. E.; Alvarez, D.; Skelly, A.; von Andrian, U. H.; Kasper, D. L., Illuminating vital surface molecules of symbionts in health and disease. Nat Microbiol 2017, 2, 17099.

32. Pidgeon, S. E.; Pires, M. M., Cell Wall Remodeling of Staphylococcus aureus in Live Caenorhabditis elegans. Bioconjug Chem 2017, 28 (9), 2310–2315.

33. Fura, J. M.; Sabulski, M. J.; Pires, M. M., D-amino acid mediated recruitment of endogenous antibodies to bacterial surfaces. ACS Chem Biol 2014, 9 (7), 1480–9.

34. Fura, J. M.; Kearns, D.; Pires, M. M., D-Amino Acid Probes for Penicillin Binding Protein-based Bacterial Surface Labeling. J Biol Chem 2015, 290 (51), 30540–50.

35. Blackman, M. L.; Royzen, M.; Fox, J. M., Tetrazine ligation: fast bioconjugation based on inverse-electron-demand Diels-Alder reactivity. J Am Chem Soc 2008, 130 (41), 13518–9.

36. Wu, K.; Yee, N. A.; Srinivasan, S.; Mahmoodi, A.; Zakharian, M.; Mejia Oneto, J. M.; Royzen, M., Click activated protodrugs against cancer increase the therapeutic potential of chemotherapy through local capture and activation. Chem Sci 2021, 12 (4), 1259–1271.

37. Korem, T.; Zeevi, D.; Suez, J.; Weinberger, A.; Avnit-Sagi, T.; Pompan-Lotan, M.; Matot, E.; Jona, G.; Harmelin, A.; Cohen, N.; Sirota-Madi, A.; Thaiss, C. A.; Pevsner-Fischer, M.; Sorek, R.; Xavier, R.; Elinav, E.; Segal, E., Growth dynamics of gut microbiota in health and disease inferred from single metagenomic samples. Science 2015, 349 (6252), 1101–1106.

38. Myhrvold, C.; Kotula, J. W.; Hicks, W. M.; Conway, N. J.; Silver, P. A., A distributed cell division counter reveals growth dynamics in the gut microbiota. Nat Commun 2015, 6, 10039.

39. Hill, D.; Sugrue, I.; Tobin, C.; Hill, C.; Stanton, C.; Ross, R. P., The Lactobacillus casei Group: History and Health Related Applications. Front Microbiol 2018, 9, 2107.

40. Human Microbiome Project, C., Structure, function and diversity of the healthy human microbiome. Nature 2012, 486 (7402), 207–14.

41. Pidgeon, S. E.; Apostolos, A. J.; Nelson, J. M.; Shaku, M.; Rimal, B.; Islam, M. N.; Crick, D. C.; Kim, S. J.; Pavelka, M. S.; Kana, B. D.; Pires, M. M., L,D-Transpeptidase Specific Probe Reveals Spatial Activity of Peptidoglycan Cross-Linking. ACS Chem Biol 2019, 14 (10), 2185–2196.

42. Dalesandro, B. E.; Pires, M. M., Induction of Endogenous Antibody Recruitment to the Surface of the Pathogen Enterococcus faecium. ACS Infect Dis 2021, 7 (5), 1116–1125.

43. Apostolos, A. J.; Pidgeon, S. E.; Pires, M. M., Remodeling of Cross-bridges Controls Peptidoglycan Cross-linking Levels in Bacterial Cell Walls. ACS Chem Biol 2020, 15 (5), 1261–1267.

44. Apostolos, A. J.; Nelson, J. M.; Silva, J. R. A.; Lameira, J.; Achimovich, A. M.; Gahlmann, A.; Alves, C. N.; Pires, M. M., Facile Synthesis and Metabolic Incorporation of m-DAP Bioisosteres Into Cell Walls of Live Bacteria. ACS Chem Biol 2020, 15 (11), 2966–2975.

45. Ngadjeua, F.; Braud, E.; Saidjalolov, S.; Iannazzo, L.; Schnappinger, D.; Ehrt, S.; Hugonnet, J. E.; Mengin-Lecreulx, D.; Patin, D.; Etheve-Quelquejeu, M.; Fonvielle, M.; Arthur, M., Critical Impact of Peptidoglycan Precursor Amidation on the Activity of l,d-Transpeptidases from Enterococcus faecium and Mycobacterium tuberculosis. Chemistry 2018, 24 (22), 5743–5747.

46. Gautam, S.; Kim, T.; Shoda, T.; Sen, S.; Deep, D.; Luthra, R.; Ferreira, M. T.; Pinho, M. G.; Spiegel, D. A., An Activity-Based Probe for Studying Crosslinking in Live Bacteria. Angew Chem Int Ed Engl 2015, 54 (36), 10492–6.

47. Lin, H.; Lin, L.; Du, Y.; Gao, J.; Yang, C.; Wang, W., Biodistributions of l,d-Transpeptidases in Gut Microbiota Revealed by In Vivo Labeling with Peptidoglycan Analogs. ACS Chem Biol 2021, 16 (7), 1164–1171.

48. Pidgeon, S. E.; Apostolos, A. J.; Nelson, J. M.; Shaku, M.; Rimal, B.; Islam, M. N.; Crick, D. C.; Kim, S. J.; Pavelka, M. S.; Kana, B. D.; Pires, M. M., L,D-Transpeptidase Specific Probe Reveals Spatial Activity of Peptidoglycan Cross-Linking. ACS Chemical Biology 2019, 14 (10), 2185–2196.

49. Turnbaugh, P. J.; Ridaura, V. K.; Faith, J. J.; Rey, F. E.; Knight, R.; Gordon, J. I., The effect of diet on the human gut microbiome: a metagenomic analysis in humanized gnotobiotic mice. Sci Transl Med 2009, 1 (6), 6ra14.

50. Stefka, A. T.; Feehley, T.; Tripathi, P.; Qiu, J.; McCoy, K.; Mazmanian, S. K.; Tjota, M. Y.; Seo, G. Y.; Cao, S.; Theriault, B. R.; Antonopoulos, D. A.; Zhou, L.; Chang, E. B.; Fu, Y. X.; Nagler, C. R., Commensal bacteria protect against food allergen sensitization. Proc Natl Acad Sci U S A 2014, 111 (36), 13145–50.

