## Supporting Information for "Non-invasive Fluorescence Imaging of Gut Commensal Bacteria in Live Mice"

`

`

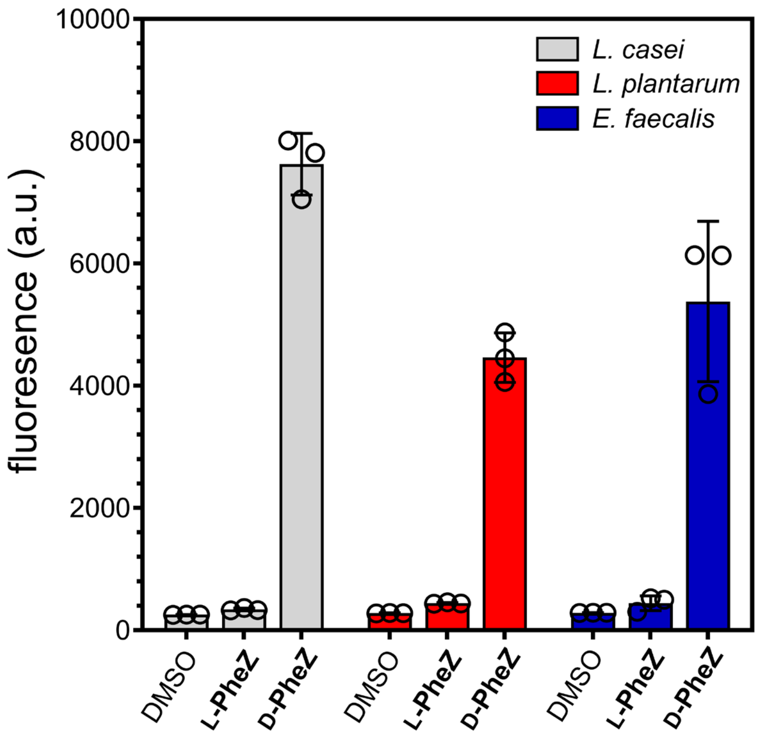


**Figure S1.** Flow cytometry analysis of *L. casei*, *L. plantarum*, and *E. faecalis* grown overnight with 25 μM **D-** or **L-PheZ** followed by treatment with 25 μM TCO-Cy5. Data are represented as mean +/- SD (n = 3). *P*-values were determined by a two-tailed *t*-test (* denotes a *p*-value < 0.05, ** < 0.01, ***<0.001, ns = not significant).

`

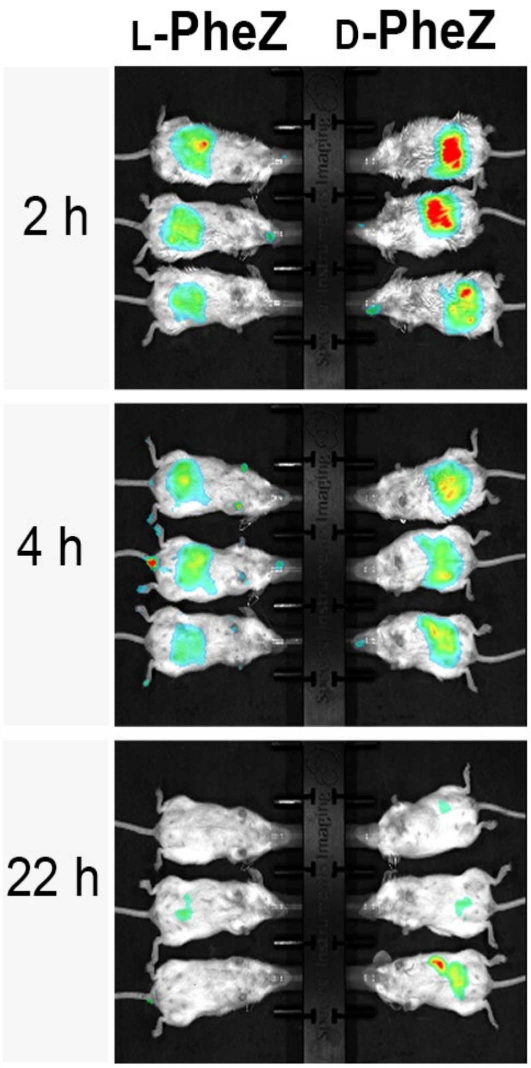


**Figure S2.** IVIS imaging of WT female mice orally gavaged with **D-PheZ** or **L-PheZ** (5 mM in 200 μL of PBS) twice (1 hour apart). After 4 hours, mice were orally administered TCO-Cy7.5 (1 mM in 100 μL of PBS). Imaging was performed 2 hours, 4 hours, and 22 hours after dye administration.

`

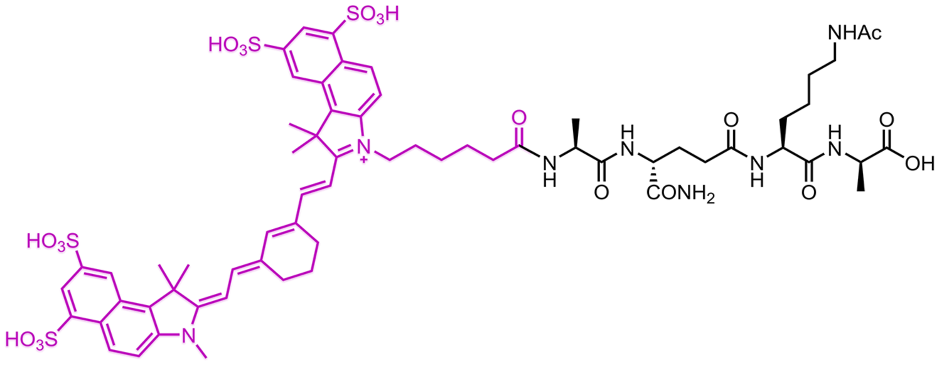


**D-Tet**

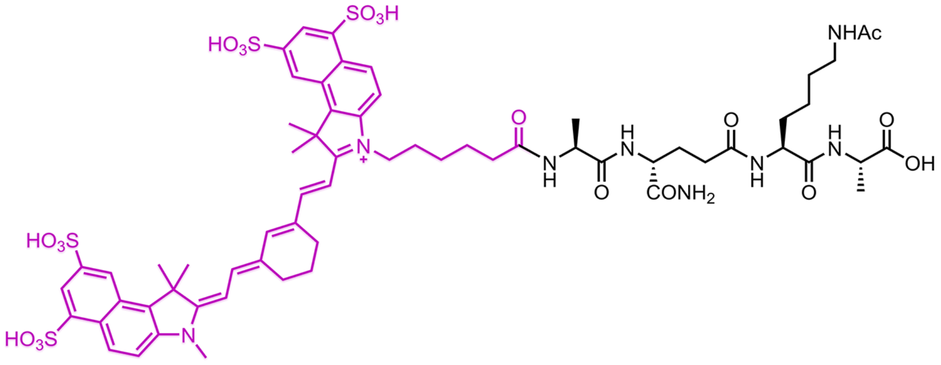


**L-Tet**

**Figure S3.** Chemical structure of **D-Tet** and **L-Tet**.

`

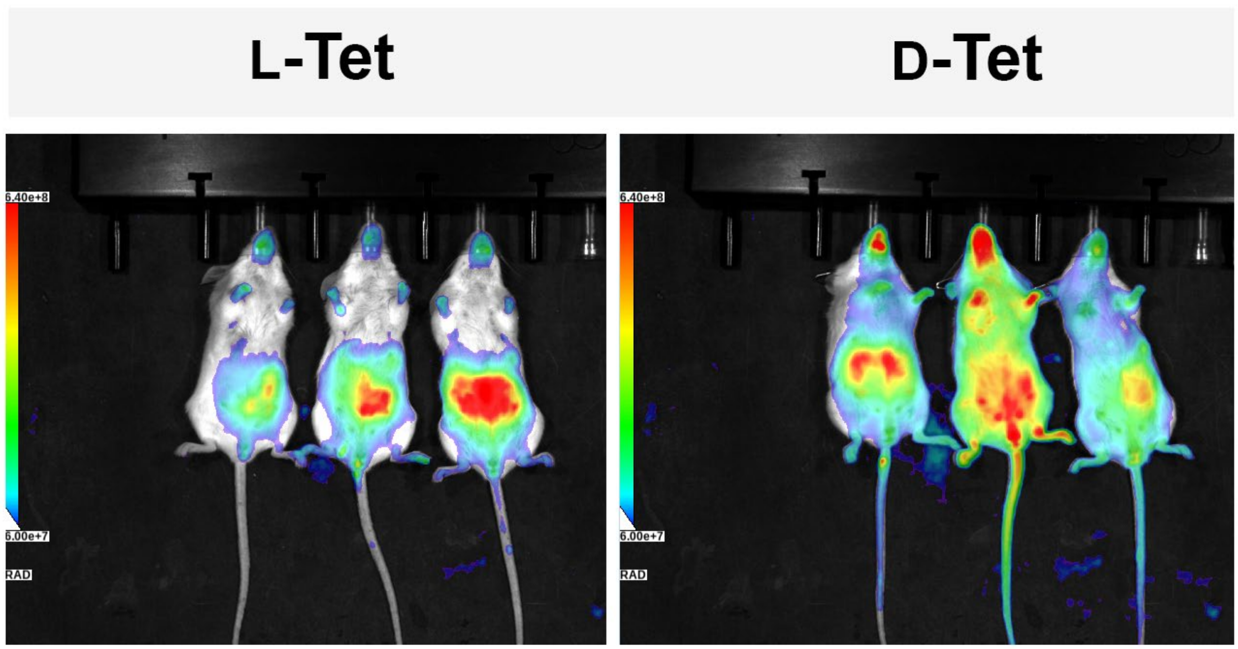


**Figure S4.** IVIS imaging of female mice orally gavaged 2x, 1 h apart, with a tetrapeptide tag (1 mM in 100 μL of PBS). Mice were imaged 4 hours after the final administration.

`

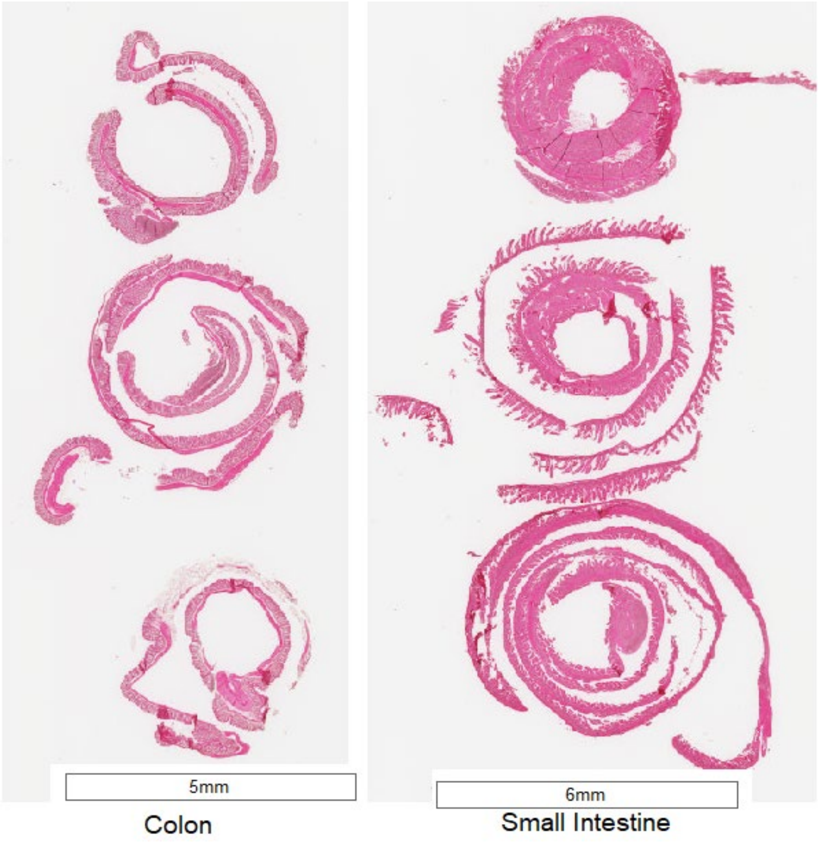


**Figure S5.** Swiss rolls were prepared from colons (left panel) and small intestines (right panel) of mice treated with **D-Tet** 22 hours after mice were last gavaged with the imaging probe.

`

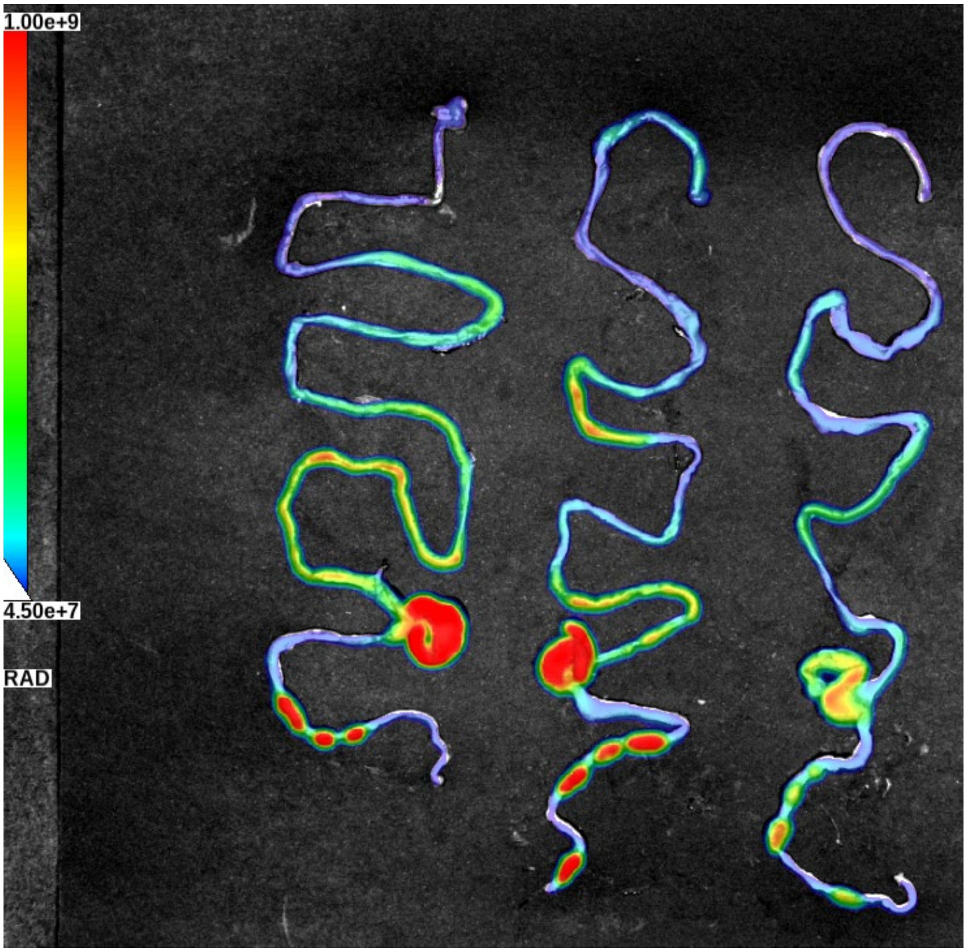


**Figure S6.** Small and large intestines harvested from GF mice that were conventionalized by FMT and then gavaged with 100 uL of 1 mM **D-Tet** twice, one hour apart. Mice were sacrificed 22 h post final gavage and intestines were imaged using LagoX IVIS (ex 745 nm, em 810 nm, 30s exposure).

`

**Materials.** All peptide related reagents (resin, coupling reagent, deprotection reagent, amino acids, and cleavage reagents) were purchased from ChemImpex. Sulfo-Cy7.5-NHS was purchase from Lumiprobe. TCO-Amine was purchased from Click Chemistry Tools. Nickel (II) trifluoromethanesulfonate, anhydrous hydrazine, and sodium nitrite were purchased from Sigma Aldrich. Bacterial strains *L. casei* and *L. plantarum* were grown in lactobacillus MRS broth for *in vitro* studies. *E. faecium* 29212 was grown in Brain Heart Infusion (BHI) broth for *in vitro* studies. The mice in this experiment were handled and processed in accordance with the protocol approved by the University of Virginia Animal Care and Use Committee in accordance with current guidelines of National Institutes of Health Model Procedure of Animal Care and Use.

***In vitro* flow cytometry analysis of bacterial labeling with single amino acid probes.** Media containing 25 µM of each single amino acid were prepared. Bacterial cells from an overnight culture were added to the medium (1:100 dilution) and allowed to grow overnight at 37°C with shaking at 250 rpm. The bacteria were harvested at 6,000g and washed three times with original culture volume of 1X PBS followed by fixation with 2% formaldehyde in 1X PBS for 30 min at room temperature. The cells were washed once more to remove formaldehyde and then resuspended in 50 µM of Cy5-TCO (Click Chemistry Tools) for 30 min at room temperature with shaking. Cells were washed three times with 1X PBS before analysis with an AttuneNxT (Thermo Fisher) flow cytometer equipped with a 638 nm laser and 670/14 nm bandpass filter. The data were analyzed using the AttuneNxT Software, where populations were gated and no less than 10,000 events per sample were recorded.

***In vitro* flow cytometry analysis of bacterial labeling with tetrapeptide probes.** Media containing 100 µM of each probe were prepared. Bacterial cells from an overnight culture were added to the medium (1:100 dilution) and allowed to grow overnight at 37°C with shaking at 250 rpm. The bacteria were harvested at 6,000g and washed three times with original culture volume of 1X PBS followed by fixation with 2% formaldehyde in 1X PBS for 30 min at room temperature. The cells were washed once more to remove formaldehyde and then analyzed using a Cytoflex S (Beckmann Coulter) flow cytometer equipped with an 808 nm laser and 840/20 nm bandpass filter. The data were analyzed using the CytExpert Software, where populations were gated and no less than 10,000 events per sample were recorded.

**Mice.** 15 – 16 week-old female BALB/c mice were purchased from Charles River laboratories and maintained in pathogen-free barrier facilities at the University of Virginia. Germ-free 16 week-old female BALB/c mice were purchased from Taconic laboratories and gavaged/imaged immediately upon arrival to UVA. Following the first round of imaging, germ-free mice were conventionalized by fecal oral gavage as previously reported (cite PMID 31064848). Briefly, cecal contents from wild-type mice housed in pathogen-free conditions at UVA were collected, homogenized, and frozen at -80C in sterile 1:1 glycerol/PBS. Germ-free mice were orally gavaged for three consecutive days with the cecal slurry, and allowed to engraft for two weeks. During this period, and prior to the second round of imaging, conventionalized germ-free mice were co-housed with wild-type mice. All experiments in this study were approved by the University of Virginia Institutional Animal Care and Use Committee.

***In vivo* gut labeling and imaging of mice with single amino acid probes (two step).** Mice were orally gavaged twice with 200 µL of 5 mM of **D-PheZ** or **L-PheZ**, one hour apart. Four hours later, mice were orally gavaged with 100 µL of 1 mM Sulfo-Cy7.5-TCO. Images of live mice were taken at selected time points after the final gavage. To image, mice were anesthetized with vaporized isoflurane (covetrus) by use of an anesthetic machine to enable imaging. Images were taken using Spectral Instruments Lago X bioluminescence, fluorescence,

`

and X-ray scanner with the fluorescence excitation set to 745 nm and the emission set to 810 nm. Images were processed using the Spectral Imaging software.

***In vivo* gut labeling and imaging of mice with tetrapeptide probes (one step).** Mice were orally gavaged with 100 µL of 1 mM of each probe twice, one hour apart. Images were taken at selected time points starting two hours after the final gavage. To image, mice were anesthetized and imaged with the parameters previously described.

***In vivo* gut labeling and imaging of GF mice with tetrapeptide probes (one step).** Mice (BALB/cAnNTac from Tacomic) were orally gavaged with 100 µL of 1 mM of each probe twice, one hour apart. Images of live mice were taken at selected time points starting two hours after the final gavage, using the parameters previously described. Conventionalized mice were orally gavaged with 100 µL of 1 mM of each probe twice, one hour apart. Images of live mice were taken at selected time points starting two hours after the final gavage, followed by imaging as described above.

`

**Scheme S1. Synthesis of D- and L-Phe-Z.**

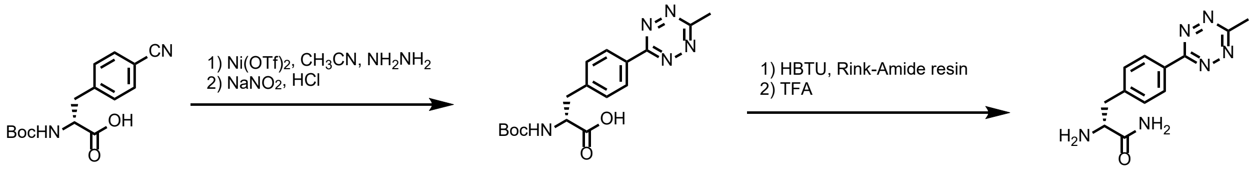


Boc-4-cyano-D-phenylalanine (1.00 g, 3.45 mmol) or Boc-4-cyano-L-phenylalanine were added to a reaction vessel with a stir bar. Nickel (II) trifluoromethanesulfonate (61.6 mg, 0.17 mmol), acetonitrile (1.80 mL, 34.5 mmol), and anhydrous hydrazine (5.50 mL, 172 mmol) were added. The vessel was sealed, and the mixture stirred in an oil bath for 24 hr at 60ºC. The solution was cooled to room temperature, followed by the dropwise addition of sodium nitrite (4.76g, 69.0 mmol) in 8 mL of water. 1M HCl was added dropwise until gas evolution ceased and pH of the solution was 3. The mixture was extracted with EtOAc, organic phase dried with $MgSO_4$. The Boc-protected tetrazine was dissolved in DMF (20 mL), followed by the addition of HBTU (3 eq, 3.16g, 8.33 mmol) and DIEA (6 eq, 2.90 mL, 16.6 mmol). The mixture was added to a peptide vessel containing rink amide resin (5.00g, 2.05 mmol) and the vessel was incubated for 2 hr at ambient temperature with shaking. The resin was washed with DCM, MeOH, DCM, MeOH, DCM 2X (15 mL each). The resin was transferred to a round bottom flask and a solution of TFA/DCM (30:70, 30 mL) was added and stirred for 1 hr on ice. The residue was triturated with cold diethyl ether and the precipitate was purified by reversed phase HPLC (RP-HPLC) using a C18(2) column (Phenomenex) to yield **D-PheZ** or **L-PheZ** as a pink solid.

`

Analytical HPLC analysis of **D-PheZ**. Solvent A (H2O 0.01% TFA) Solvent B (MeCN 0.01% TFA). 5% Solvent B to 100% Solvent B over 50 mins; absorbance measured at 220 nm.

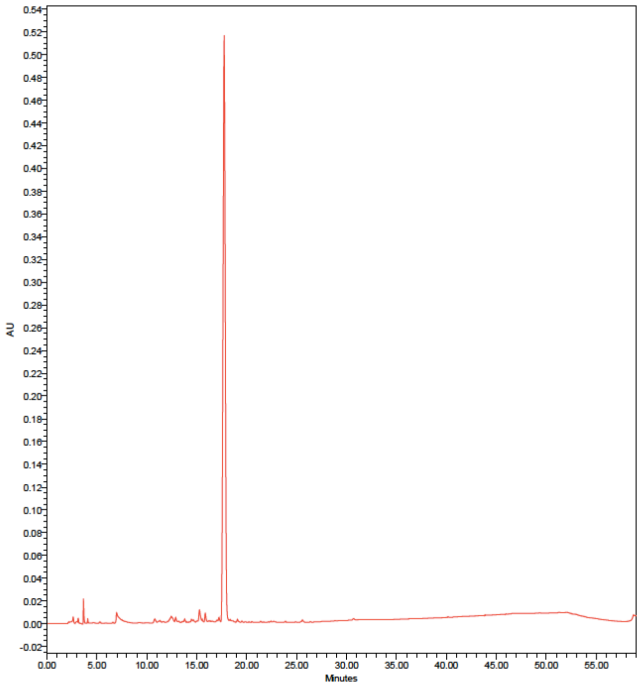

`

M+H⁺ calculated = 259.1302, observed = 259.1302

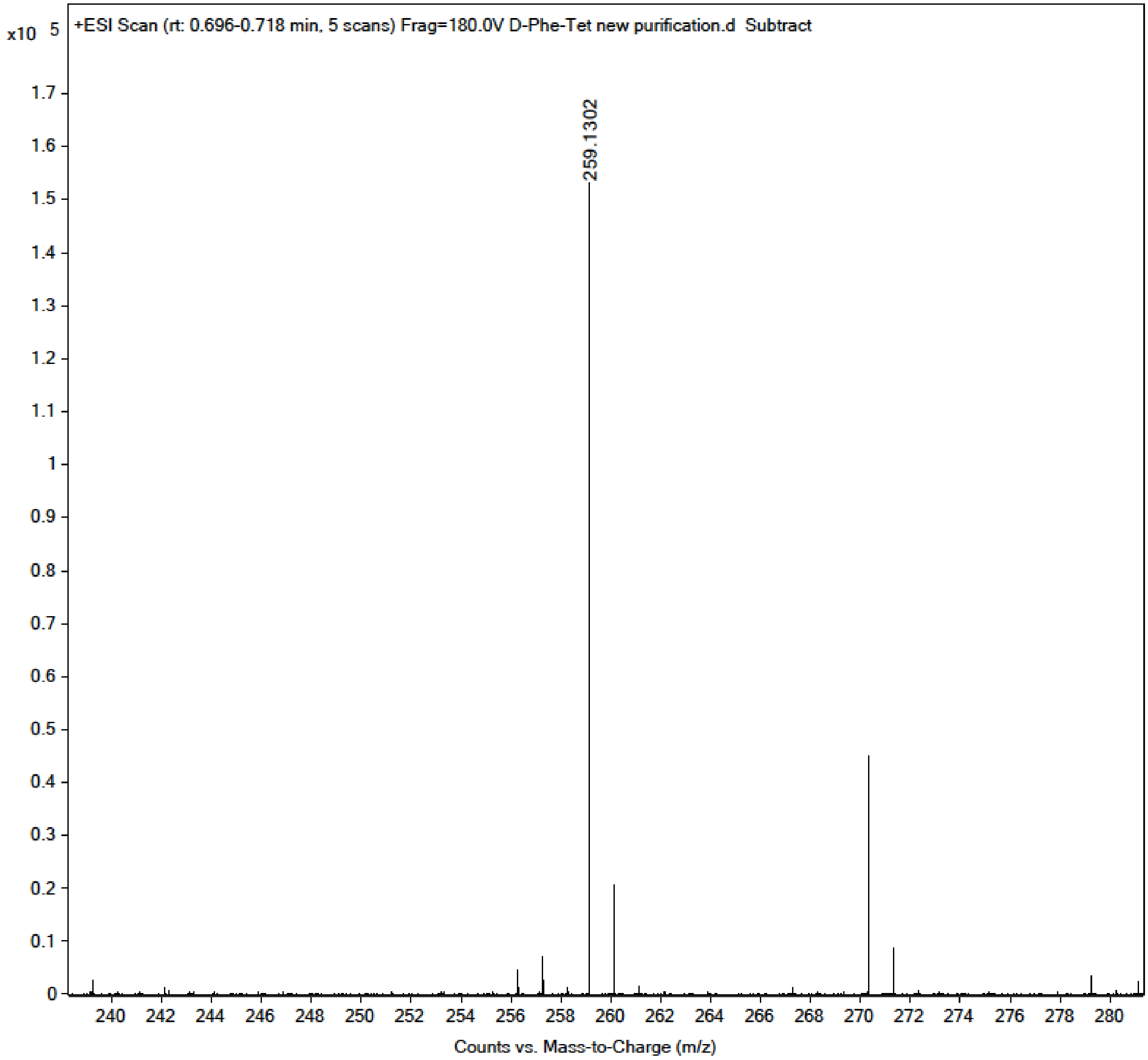

`

[1]H NMR (600 MHz, $CD_3OD$) δ 8.56 (d, *J* = 6.0 Hz, 2H, 2'-ArH), 7.58 (d, *J* = 6.0 Hz, 2H, 3'-ArH), 4.19 (t, J=6.0 Hz, 1H, CH), 3.37 (dd, J=6.0, 18Hz, 1H, CH2), 3.19 (dd, J=6.0, 18Hz, 1H, CH2), 3.05 (s, 3H, CH3).

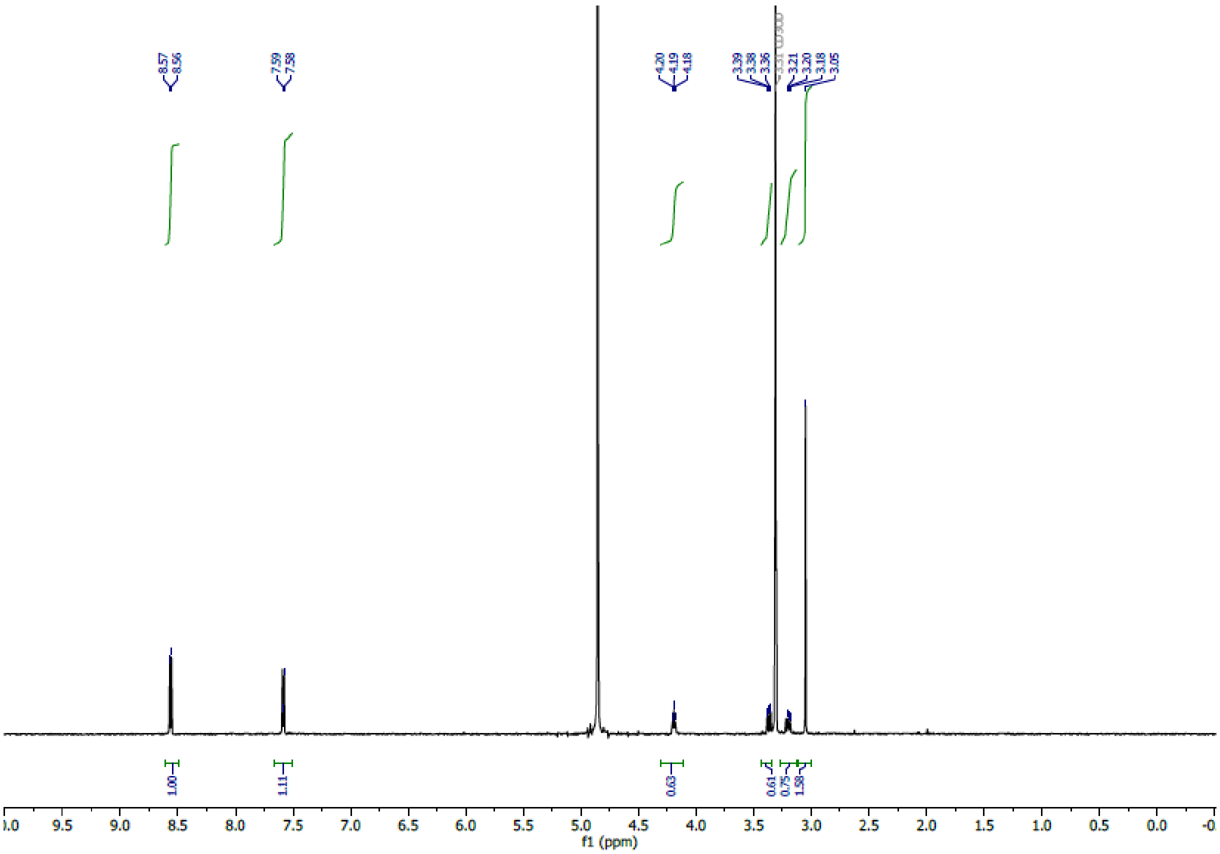

`

Analytical HPLC analysis of **L-PheZ**. Solvent A (H2O 0.01% TFA) Solvent B (MeCN 0.01% TFA). 5% Solvent B to 100% Solvent B over 50 mins; absorbance measured at 220 nm.

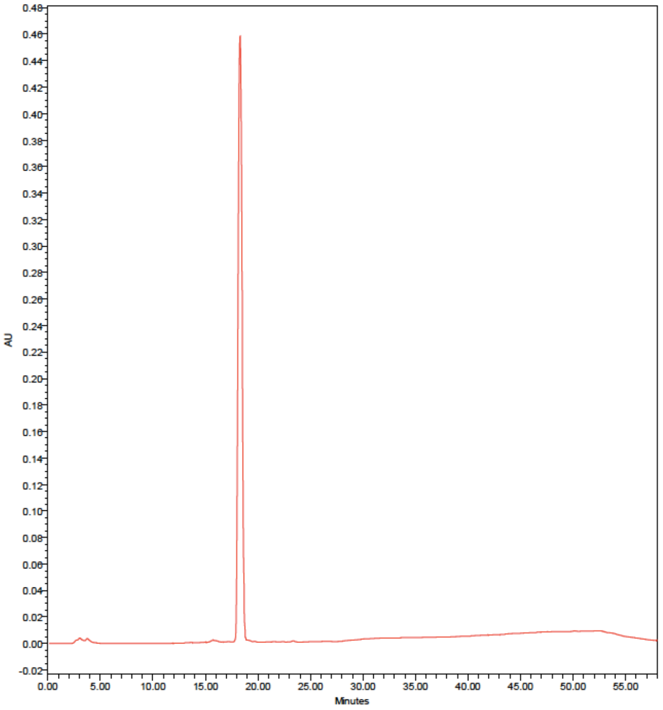

`

M+H$^+$ calculated = 259.1302, observed = 259.1297
M+Na$^+$ calculated = 281.1121, observed = 281.1117

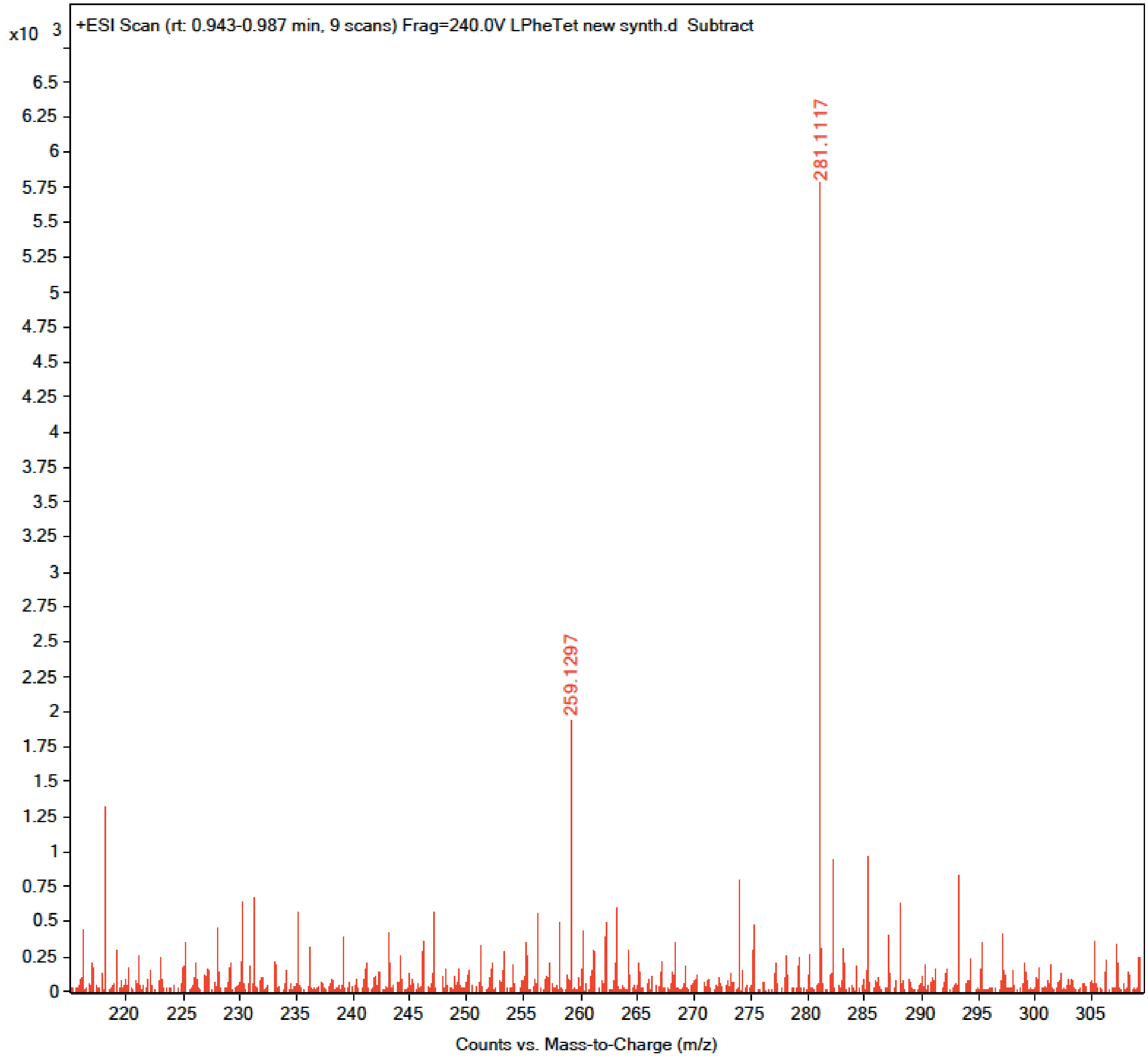

`

$^{1}H$ NMR (600 MHz, $CD_3OD$) δ 8.56 (d, *J* = 6.0 Hz, 2H, 2'-ArH), 7.58 (d, *J* = 6.0 Hz, 2H, 3'-ArH), 4.18 (t, J=6.0 Hz, 1H, CH), 3.37 (dd, J=6.0, 18Hz, 1H, CH2), 3.20 (dd, J=6.0, 18Hz, 1H, CH2), 3.05 (s, 3H, CH3).

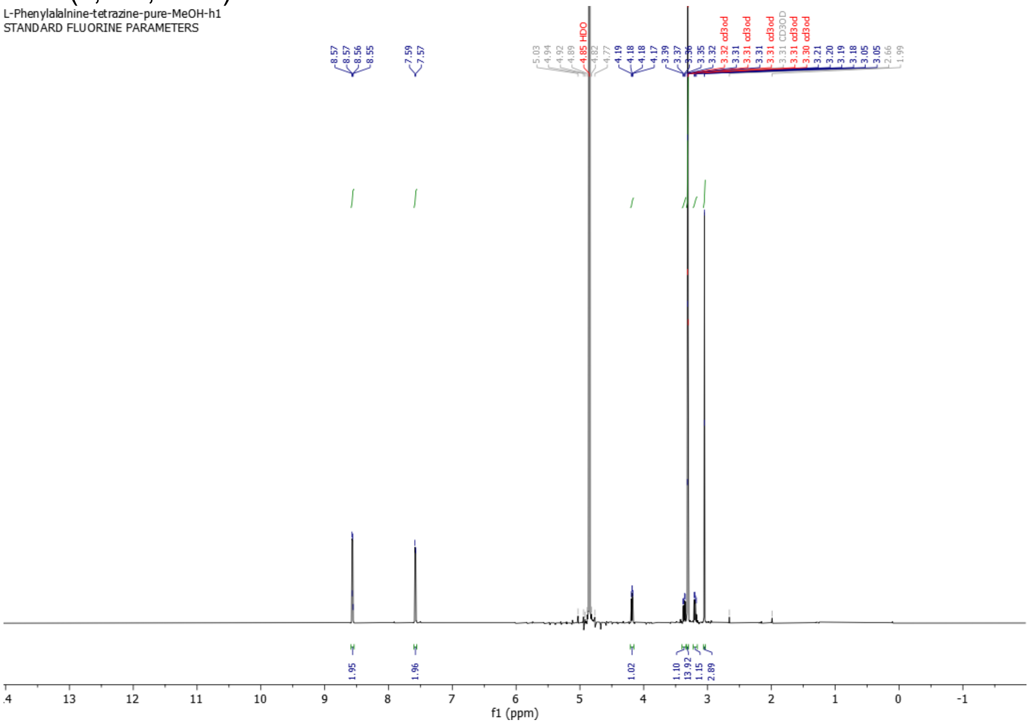

`

**Scheme S2. Synthesis of SulfoCy7.5-TCO**

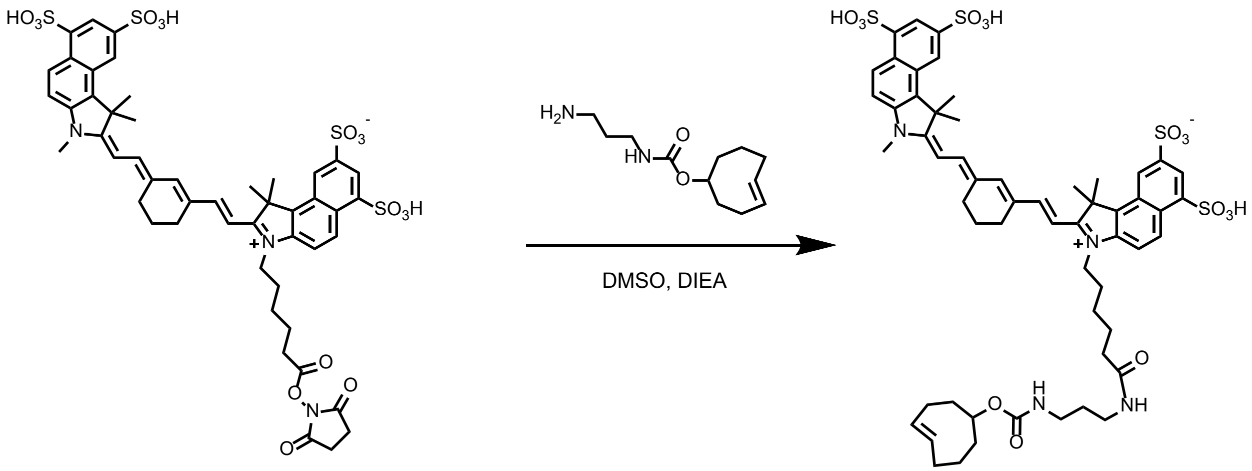


SulfoCy7.5 NHS Ester (Lumiprobe) was reacted with TCO-Amine HCl (Click Chemistry Tools) and DIEA was added to the solution (in DMSO, 1 Sulfo Cy7.5 NHS Ester : 1 TCO-Amine : 7 DIEA). The reaction was purified using RP-HPLC equipped with a C8(2) column.

Analytical HPLC analysis of **SulfoCy7.5-TCO**. Solvent A (H2O 0.01% TFA) Solvent B (MeCN 0.01% TFA). 5% Solvent B to 100% Solvent B over 50 mins; absorbance measured at 220 nm.

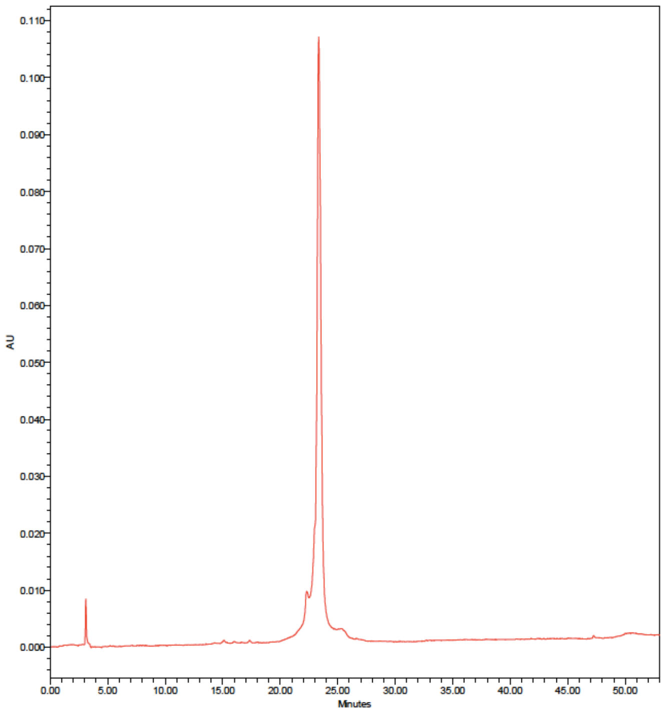

`

M+H⁺ calculated = 1178.3710, observed = 1178.7711
M+2H⁺ calculated = 589.6891, observed = 589. 4264

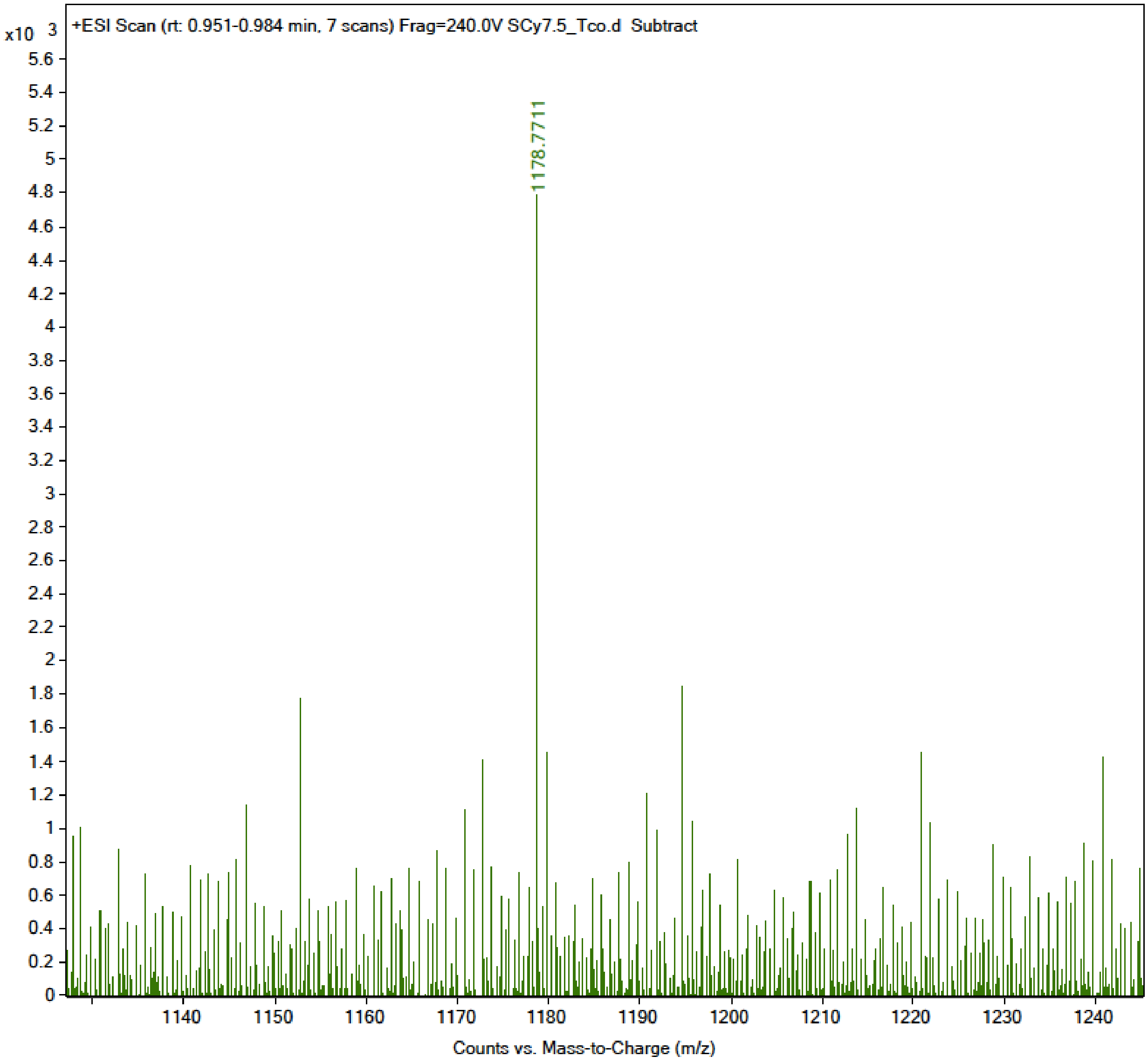

`

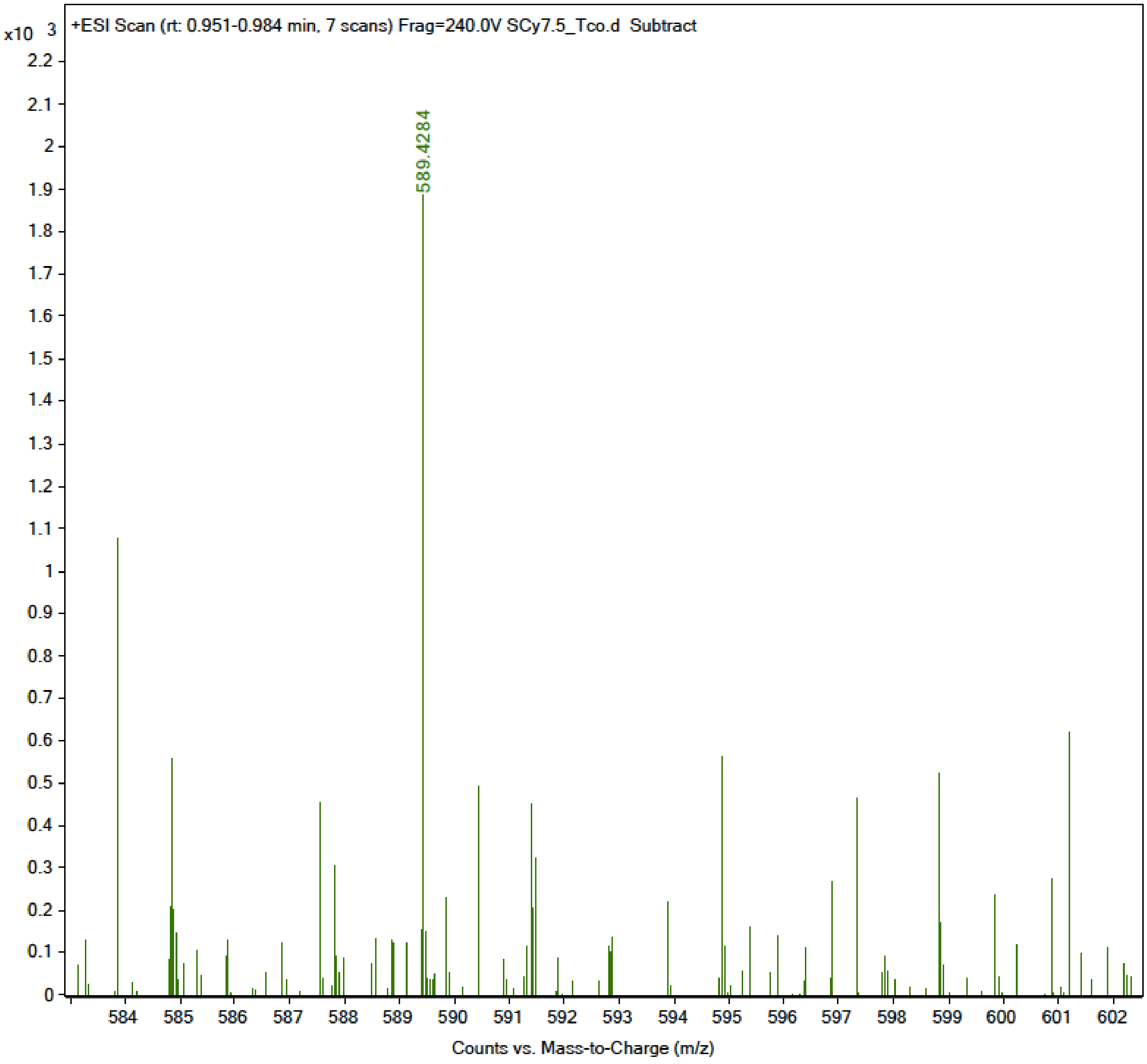
+ESI Scan (rt: 0.951-0.984 min, 7 scans) Frag=240.0V SCy7.5_Tco.d Subtract
x10 3
589.4284
Counts vs. Mass-to-Charge (m/z)

`

$^{1}$H NMR (600 MHz, $CD_3OD$)

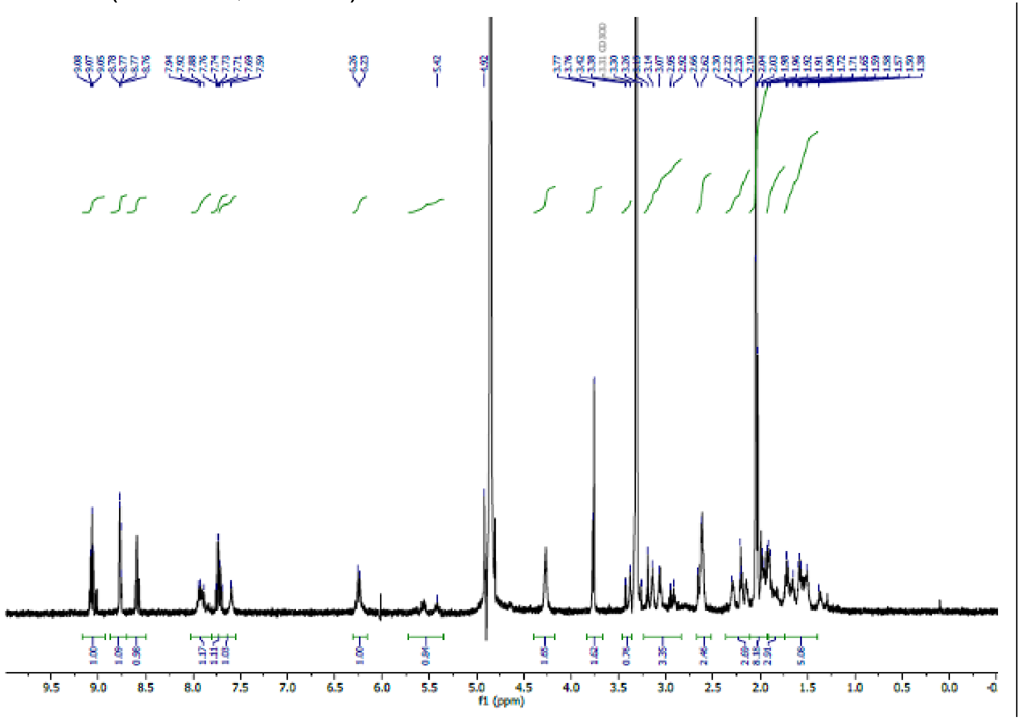

`

**Scheme S3. Synthesis of D- or L-Tet**

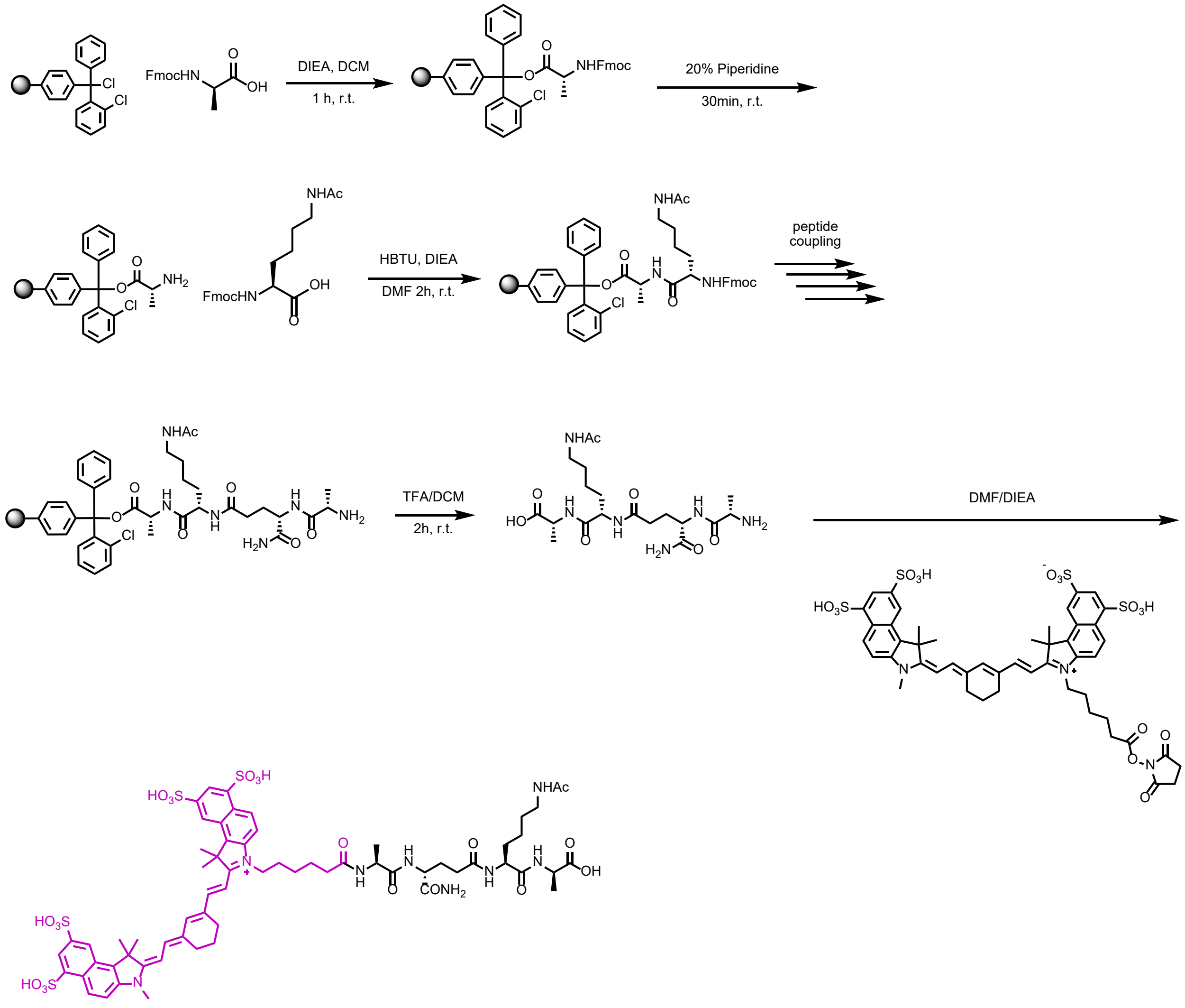


Fmoc-D-Ala-OH **or** Fmoc-L-Ala-OH (1.1 eq, 188 mg, 0.605 mmol) was added to a 25 mL peptide synthesis vessel charged with 2-chlorotrityl chloride resin (500 mg, 0.55 mmol) and DIEA (4 eq, 0.382 mL, 2.20 mmol) in dry DCM (5 mL). The resin was agitated for 1 h at ambient temperature and washed with MeOH and DCM (3 x 15 mL each). The Fmoc protecting group was removed with 20% piperidine in DMF (15 ml) for 30 min at ambient temperature, then washed as before. Fmoc-L-Lys(Ac)-OH (3 eq, 677 mg, 1.65 mmol), HBTU (3 eq, 625 mg, 1.65 mmol), and DIEA (6 eq, 0.574 mL, 3.30 mmol) in DMF (10 mL) were added to the reaction flask and agitated for 2 h at ambient temperature. The Fmoc deprotection and coupling procedure was repeated as before using the same equivalencies with Fmoc-D-glutamic acid α-amide (3 eq, 608 mg, 1.65 mmol) and then with Fmoc-L-Alanine-OH (3 eq, 514 mg, 1.65 mmol). The Fmoc group at the *N*-terminus was removed and the peptide was cleaved from resin with a solution of TFA/$H_2O$/TIPS (95%, 2.5%, 2.5%, 20 mL) with agitation for 2 h at ambient temperature. The resin was filtered and resulting solution concentrated *in vacuo*. The residue was triturated with cold diethyl ether and then reacted with Sulfo-Cy7.5 NHS ester (2 mg) in DMF (250 µL) and DIEA (50 µL, 0.288 mmol) for 4 h at room temperature. The peptide was then crashed out in cold diethyl ether, concentrated *in vacuo*, and purified by RP-HPLC.

`

Analytical HPLC analysis of **L-Tet**. Solvent A (H2O 0.01% TFA) Solvent B (MeCN 0.01% TFA). 5% Solvent B to 100% Solvent B over 50 mins; absorbance measured at 220 nm.

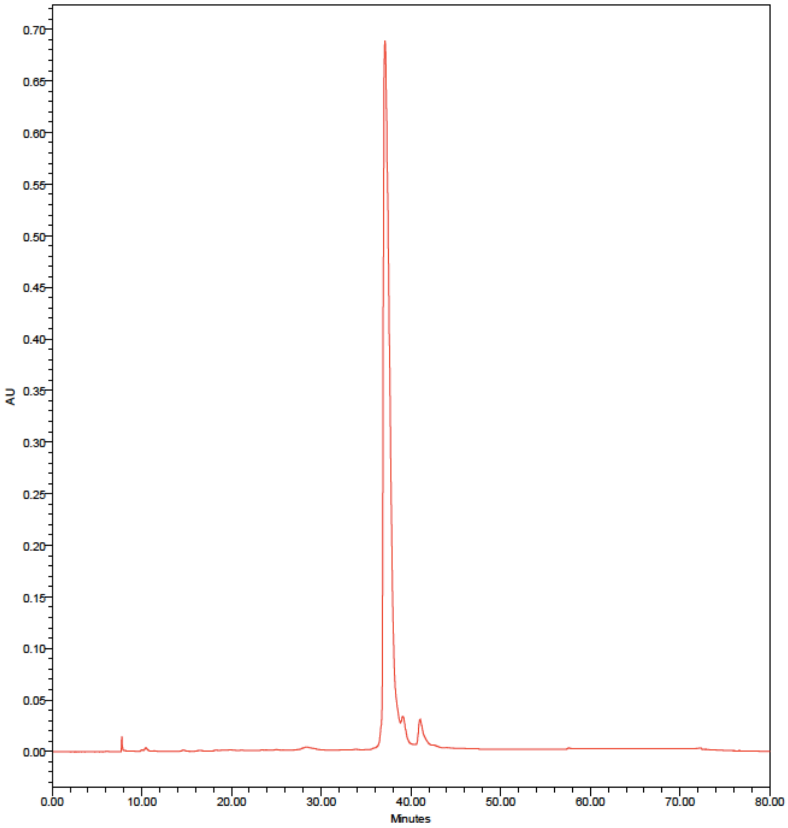

`

Expected M+H$^+$ = 1410.627, Observed = 1410.249

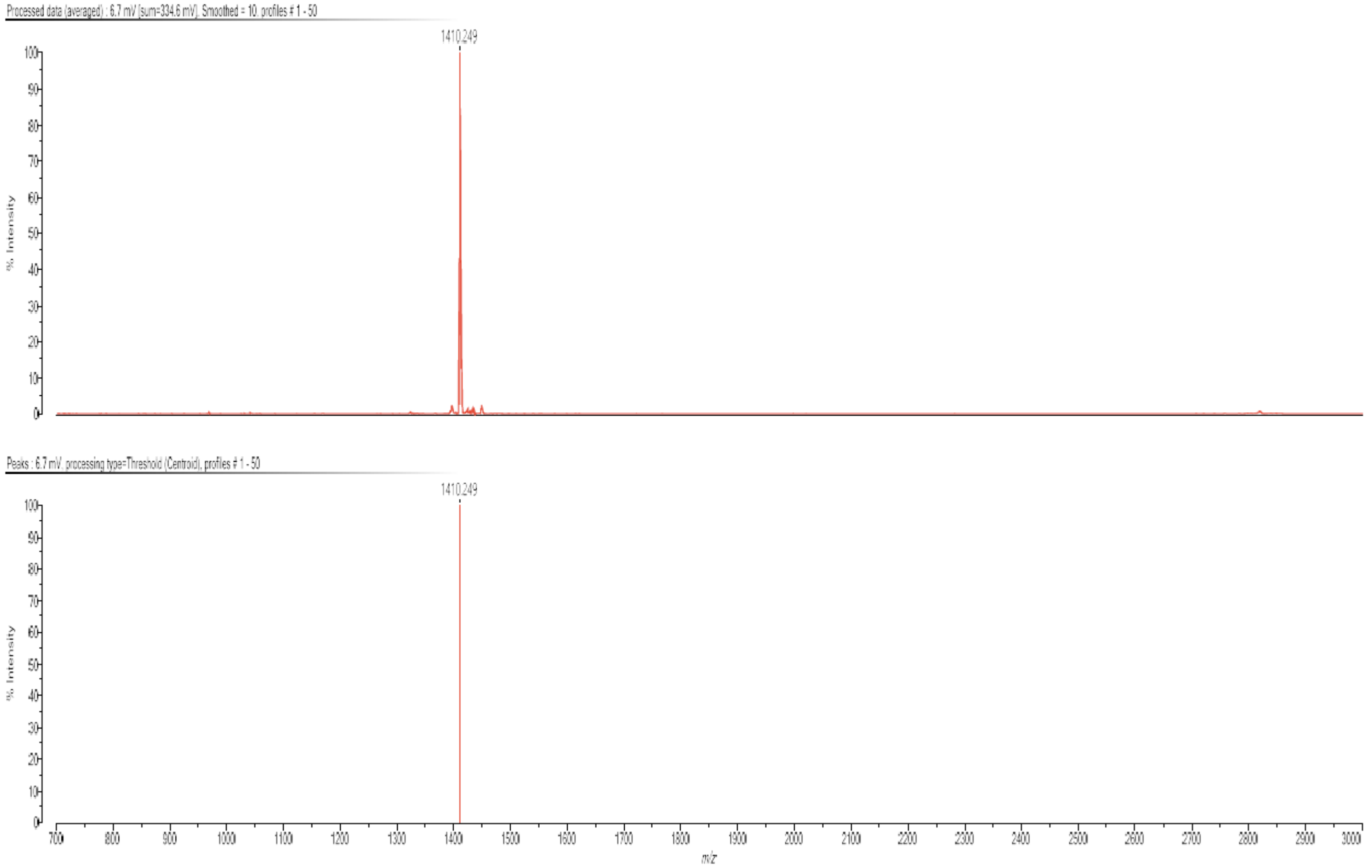

`

Analytical HPLC analysis of **D-PheZ**. Solvent A (H2O 0.01% TFA) Solvent B (MeCN 0.01% TFA). 5% Solvent B to 100% Solvent B over 50 mins; absorbance measured at 220 nm.

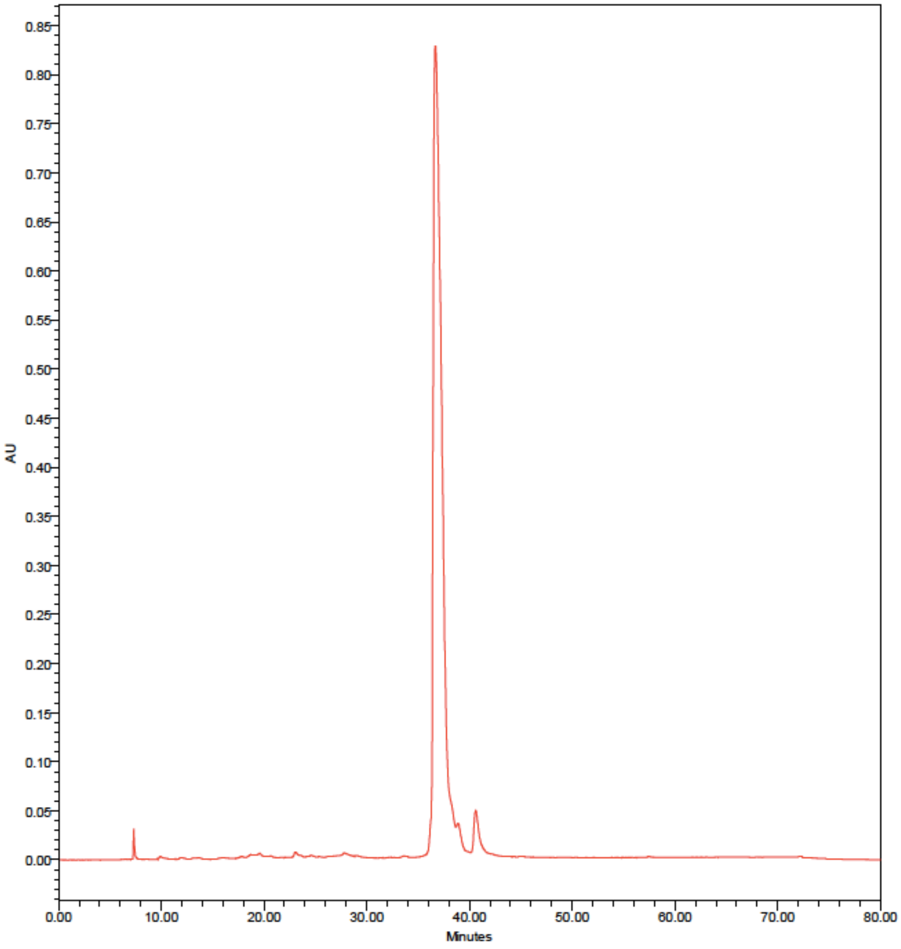

`

Expected M+H$^+$ = 1410.627, Observed = 1410.274

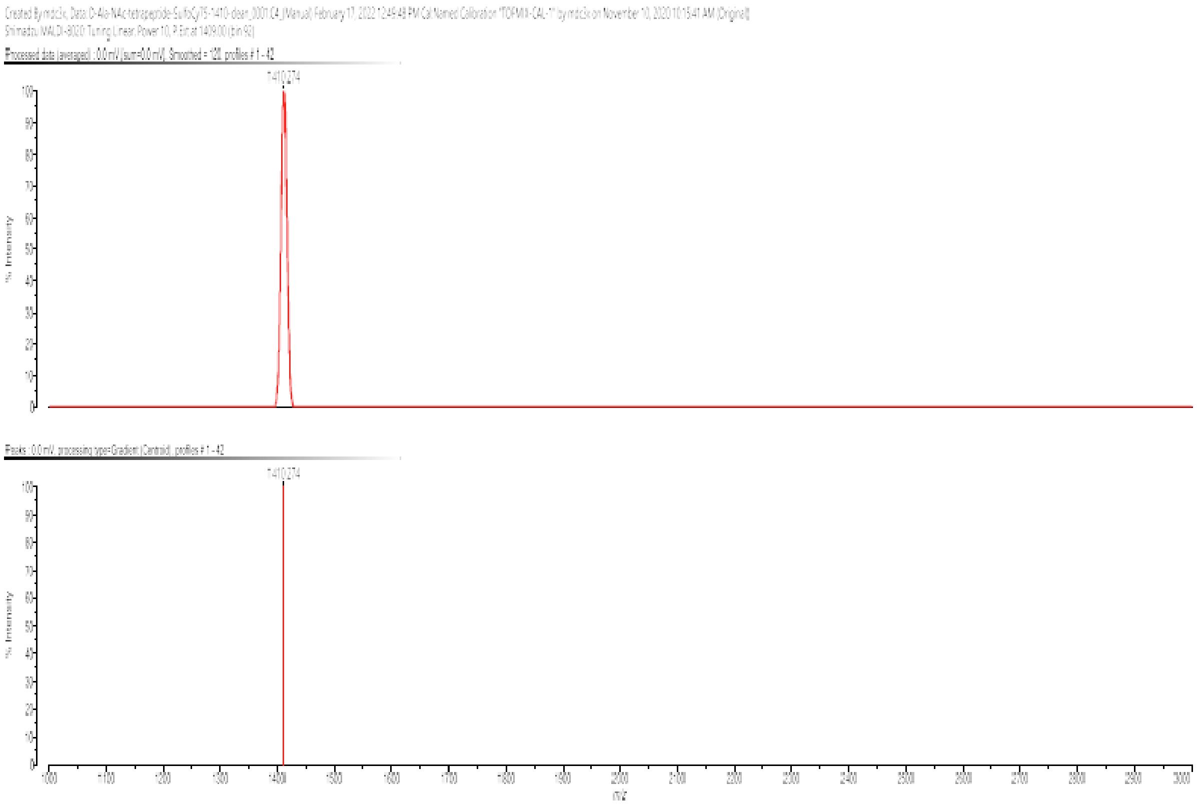

`

[1]H NMR (600 MHz, $CD_3OD$)

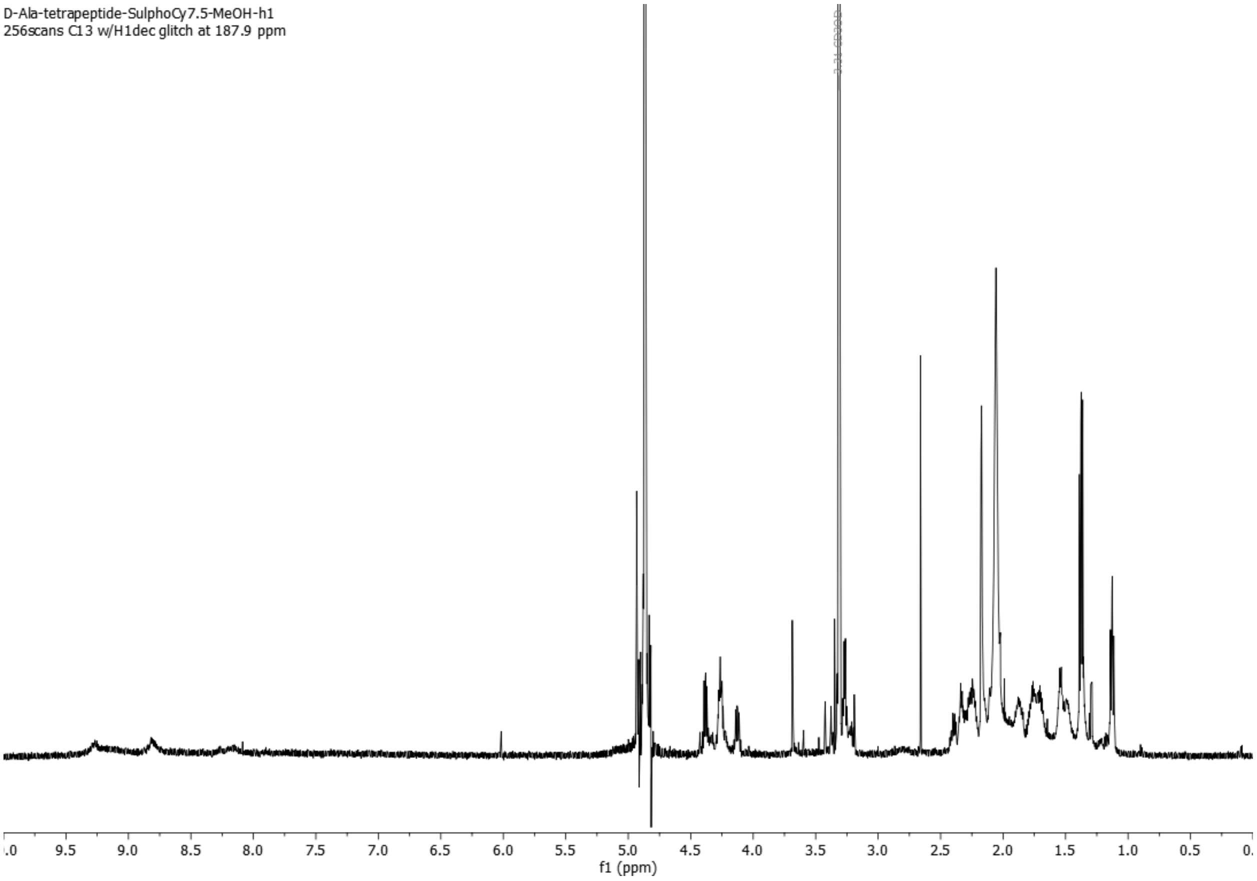
